# Cortico-thalamic dysconnectivity links with aberrant striatal dopamine in schizophrenia *A simultaneous ^18^F-DOPA-PET/resting-state fMRI study*

**DOI:** 10.1101/847178

**Authors:** Mihai Avram, Felix Brandl, Franziska Knolle, Jorge Cabello, Claudia Leucht, Martin Scherr, Mona Mustafa, Nikolaos Koutsouleris, Stefan Leucht, Sibylle Ziegler, Christian Sorg

## Abstract

In schizophrenia, among the most consistent brain changes are both aberrant dopamine function in the dorsal striatum and aberrant intrinsic functional connectivity (iFC) between distinct cortical networks and thalamic nuclei; however, it is unknown whether these changes are pathophysiologically related. Such a relationship is expected because cortico-thalamic-connectivity is modulated by striatal dopamine within topographically distinct, parallel but interacting cortico-basal-ganglia-thalamic circuits. We hypothesized: (1) Within-circuits, aberrant striatal dopamine contributes to aberrant cortico-thalamic-iFC, specifically, associative-striatum dopamine contributes to salience-network-thalamic-iFC, and sensorimotor-striatum dopamine to auditory-sensorimotor-network-thalamic-iFC. (2) Due to between-circuits interactions following an anterior-to-posterior gradient, salience-network-centered-system changes contribute to auditory-sensorimotor-network-centered-system changes. To test these hypotheses, 19 patients with schizophrenia during symptomatic remission of positive symptoms and 19 age- and sex-comparable controls underwent simultaneous fluorodihydroxyphenyl-L-alanine positron emission tomography (^18^F-DOPA-PET) and resting-state functional magnetic resonance imaging (rs-fMRI). The influx constant k_i_^cer^ based on ^18^F-DOPA-PET was used to measure dopamine synthesis capacity (DSC), indicating striatal dopamine function; correlation coefficients between rs-fMRI time-series of cortical networks and thalamic regions-of-interest were used to measure iFC. In the salience-network(SAL)-centered-system, patients had reduced associative-striatum-DSC, which correlated positively with SAL-mediodorsal-thalamus-iFC and mediated the reduction of SAL-thalamic-iFC in patients. In the auditory-sensorimotor-network(ASM)-centered-system, patients had reduced sensorimotor-striatum-DSC, which correlated positively with ASM-ventrolateral-thalamus-iFC, but did not mediate increased ASM-thalamic-iFC in patients. Finally, aberrant DSC and iFC of the SAL-centered-system mediated corresponding changes in the ASM-centered-system. Results demonstrate that cortico-thalamic-dysconnectivity links with aberrant striatal dopamine in schizophrenia - in a topographically distinct way, with an anterior-to-posterior gradient, and primary changes in the SAL-centered system.

## Introduction

Both aberrant dopamine transmission in the striatum^1-8^ and aberrant intrinsic functional connectivity (iFC) between distinct cortical networks and thalamic nuclei^9-17^ are among the most consistent findings of brain alterations in schizophrenia. As basal ganglia circuits, including the striatum and midbrain-based dopaminergic projections into the striatum, modulate cortico-thalamic-iFC, it has been suggested that these brain changes are pathophysiologically related.^9, 17-20^ The current study investigated this proposed relationship in patients with schizophrenia using simultaneous Fluorodihydroxyphenyl-L-alanine positron emission tomography (^18^F-DOPA-PET) to assess dopamine transmission and blood oxygenation level-dependent resting-state functional magnetic resonance imaging (rs-fMRI) to evaluate cortico-thalamic-iFC.

Distinct aspects of dopamine transmission have been shown to be aberrant in schizophrenia, from aberrant striatal dopamine synthesis capacity (DSC) to aberrant dopamine release and receptor availability.^1-8, 21-23^ Aberrant DSC, a presynaptic transmission measure assessed by ^18^F-DOPA-PET, is the most consistent finding in schizophrenia, particularly in the associative-striatum (i.e., mainly caudate nucleus, contrasted to the sensorimotor-striatum; i.e., mainly putamen).^5, 24^ Increased striatal-DSC is typically found in patients with significant positive symptoms^1, 4-7^ and decreased striatal-DSC during remission of positive symptoms.^8, 25^

Cortico-thalamic-dysconnectivity in schizophrenia, on the other hand, is typically indicated by aberrant iFC of rs-fMRI signals, i.e., correlated BOLD fluctuations <0.1Hz.^26^ IFC reflects coherent, infra-slow fluctuations in ongoing brain activity of distinct regions, representing basic organization of brain activity.^27-29^ Both *decreased* iFC between frontal cortices (i.e., salience-network (SAL) – covering cingulo-opercular-insular-mediofrontal cortices) and ventral anterior/mediodorsal thalamus and *increased* iFC between primary-sensorimotor cortices (i.e., auditory-sensorimotor-network (ASM) – covering lateral somatomotor-temporal cortices) and posterior/ventrolateral thalamus have been reported in at-risk, first-episode, and chronic patients with schizophrenia and first-degree relatives of patients.^10, 13,14, 17, 30, 31^

In general, anatomical studies demonstrated that the brain consists of parallel distributed networks.^32, 33^ This feature is conspicuously shown for topographically organized connections between cerebral cortex and thalamus, with integrated basal ganglia circuits.^34^ In particular, SAL-anterior-thalamus-connectivity extends to dorsomedial parts of basal ganglia circuits (including associative-striatum and corresponding midbrain dopaminergic projections) – we call it the SAL-centered-system. More posterior ASM-ventrolateral-thalamus-connections integrate dorsolateral parts of basal ganglia (including sensorimotor-striatum and corresponding midbrain dopaminergic projections) – we call it the ASM-centered-system.^17, 35-37^ A recent rs-fMRI study in healthy subjects, for example, found that SAL-iFC is modulated by striatal-DSC levels (including associative-striatum), indicating the ‘vertical’ nature of interactive processes within the SAL-centered-system.^38^ (For similar findings in humans see^19, 39^, for animal literature see^40, 41^, for a classical review see^34^, and for a more recent version see^42^) Beyond these parallel topographically organized ‘vertical’ systems, recent anatomical and physiological studies demonstrated additional ‘horizontal crosstalk’ between systems, at distinct levels and mainly with an anterior-to-posterior gradient of direction or effect.^40, 43-45^ For example, extensive within-striatum connections have been observed,^40, 46^ but also a set of indirect ventromedial-to-dorsolateral connections spiraling through the dopaminergic ventral midbrain.^43, 47^

Based on these findings, we hypothesized that (1) aberrant striatal dopamine, namely DSC, contributes to aberrant cortico-thalamic-dysconnectivity, namely iFC, within distinct parallel circuits, i.e., within SAL-centered and ASM-centered-systems, respectively (Figure 1). (2) We expected that within-SAL-centered-system alterations, namely in DSC and iFC, contribute to corresponding ASM-centered-system alterations, based on the anterior-to-posterior gradient effect described above. To test these hypotheses, we employed simultaneous ^18^F-DOPA-PET and rs-fMRI with a hybrid PET/MR scanner in healthy controls and patients with schizophrenia during symptomatic remission of positive symptoms. PET-data were derived from a sub-sample of subjects from a previous study, which demonstrated decreased striatal-DSC in schizophrenia during symptomatic remission of positive symptoms.^8^ Measures of DSC and iFC were related by partial correlation. The direct contribution of DSC to iFC was tested by mediation analysis.

**Figure 1.**
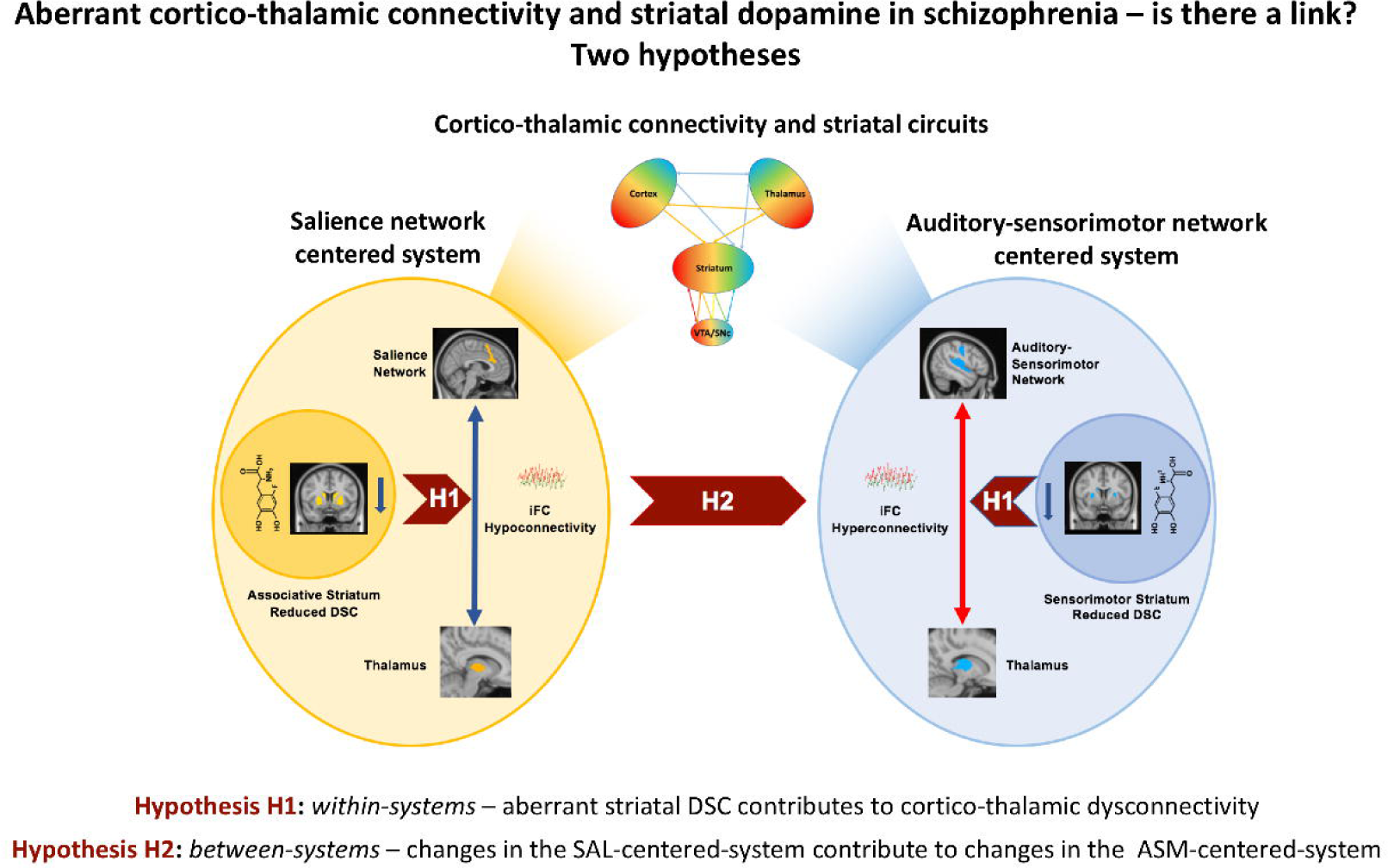
Aberrant cortico-thalamic-connectivity and striatal dopamine in schizophrenia – is there a link? Two hypotheses. *Top:* schematic depiction of cortico-thalamic-connectivity and striatal circuits embedded in larger cortico-striato-pallido-thalamo-cortical (CSPTC) circuits, following Haber and McFarland^73^. Color gradients across cortical and subcortical regions indicate topographic, parallel, but continuous organization of circuits, from which we derived anterior salience-network-centered and posterior auditory-sensorimotor-network-centered systems. Note that these circuits receive dopaminergic modulation from the VTA/SNc; particularly, striato-nigral projections form a spiral along an anterior-to-posterior gradient, which underpin – amongst other mechanisms – an anterior-to-posterior effect across CSPTC-circuits. *Left:* The salience-network-centered-system is depicted as an orange ellipse. On the left, a circle is shown containing the chemical structure of fluorodopa and a mask of the associative-striatum, together representing associative-striatum-DSC. In line with previous findings, in patients during symptomatic remission of positive symptoms, associative-striatum-DSC is reduced compared to healthy controls,^8^ depicted by a blue arrow pointing downwards. On the right side, salience-network-thalamic-iFC is depicted. On the top, a mask of the salience-network is shown^54^. On the bottom, the thalamic cluster that has been shown to be hypoconnected to the salience-network in our previous study^17^ is depicted. Hypoconnectivity between this cluster and the salience-network is depicted by a blue double-arrow. Between the associative-striatum-DSC and salience-network-thalamic-iFC, an arrow is shown containing ‘H1’, reflecting our first hypothesis. *Right*: The auditory-sensorimotor-centered-system is depicted as a blue ellipse. On the right, a circle is depicted containing the chemical structure of fluorodopa and a mask of the sensorimotor-striatum, together representing sensorimotor-striatum-DSC. Sensorimotor-striatum-DSC was also previously reported as reduced in patients during symptomatic remission of positive symptoms sensorimotor-striatum dopamine synthesis capacity,^8^ depicted by a blue arrow pointing downwards. On the left side, the auditory-sensorimotor-network-thalamic-iFC is depicted. On the top, a mask of the auditory-sensorimotor-network is shown^54^. On the bottom, the thalamic cluster that has been shown to be hyperconnected with the auditory-sensorimotor-network in our previous study^17^ is shown. Hyperconnectivity between this cluster and the auditory-sensorimotor-network is depicted by a red double-arrow. Between the sensorimotor-striatum and auditory-sensorimotor-network-thalamic-iFC, an arrow is shown containing ‘H1’, reflecting our first hypothesis. *Middle*: an arrow is depicted containing ‘H2’, reflecting our second hypothesis. *Abbreviations:* DSC – dopamine synthesis capacity, iFC – intrinsic functional connectivity, VTA - ventral tegmental area, SNc – substantia nigra pars compacta.

## Materials and Methods

### Participants

23 patients meeting DSM-IV criteria for schizophrenia (age range: 23-65 years; mean: 43.04±11.90 years) and 24 comparable healthy controls (age range: 25-62 years; mean: 38.54±11.63 years) participated in the study. Due to excessive head motion during rs-fMRI, several subjects were excluded, with 19 subjects remaining in each group for further analyses (see below). Patients had established schizophrenia but were in symptomatic remission of positive symptoms (based on severity criteria by Andreasen^48^) and were treated with antipsychotic medication (see Table S1). Substance abuse, except nicotine, was an exclusion criterion in both groups. Cognitive abilities were evaluated by the same trained rater (C.L.) for all participants with the Brief Assessment of Cognition in Schizophrenia (BACS).^49^ Participants gave written informed consent after receiving a complete description of the study, which was approved by the Ethics Review Board of the Technical University of Munich. Approval to administer radiotracers was obtained from the Administration of Radioactive Substances (Bundesamt für Strahlenschutz), Germany. (For more detailed participant description see Supplementary Methods or^8^).

### ^18^F-DOPA-PET data acquisition, preprocessing, and k_i_^cer^ as outcome measure for DSC

^18^F-DOPA-PET and MRI data were acquired simultaneously with a hybrid whole-body mMR Biograph PET/MRI scanner (Siemens-Healthineers, Erlangen, Germany). We used ^18^F-DOPA-PET to measure the influx constant k_i_^cer^, a quantitative measure reflecting DSC, in a voxel-wise manner with Gjedde–Patlak linear graphical analysis^50^, and compared averaged striatal-subdivisions’ k_i_^cer^ across groups with two-sample t-tests. The same approach was used to analyze the PET-data as in our previous study,^8^ see Supplementary Methods for more detailed description.

### MRI data acquisition, preprocessing, and correlation coefficients as outcome measure for iFC

Before rs-fMRI, participants were instructed to close their eyes, remain still, but keep awake. Rs-fMRI started simultaneously with the PET scan and lasted 20 minutes 8 seconds. 600 volumes were acquired, consisting of 35 axial slices (slice thickness=3.0mm), field of view (FOV)=192×192mm matrix (3.0mm×3.0mm in-plane resolution), 90-degree flip angle, TR=2000ms, and TE=30ms. Anatomical T1-weighted structural images were obtained using a gradient-echo scan (TR/TE/flip angle: 2.300ms/2.98ms/9°; 160 slices (gap 0.5mm) covering the whole brain; FoV: 256mm; matrix size: 256×256; voxel size: 1.0×1.0×1.0mm^3^).

MRI data were preprocessed with the Configurable Pipeline for the Analysis of Connectomes (C-PAC, http://fcp-indi.github.com). After removing the first 5 volumes, preprocessing included image realignment, motion correction, scrubbing, intensity normalization, nuisance signal regression (scanner drift, head motion signals, and component-based noise correction (Compcor; to remove physiological noise^51^), bandpass filtering (0.01-0.1 Hz), registration to anatomical space and normalization to MNI 2mm^3^ space with FSL FLIRT/FNIRT. Due to excessive head motion, estimated with mean framewise displacement (FD),^52, 53^ 5 healthy controls and 4 patients were excluded (FD>0.2mm). The remaining participants (19 per group) did not differ in mean FD (p=0.49), percentage of frames deleted (p=0.71), age (p=0.27), and injected radioligand (p=0.91) as shown by t-tests or sex (p=0.73) as shown by a chi-squared test (Table 1).

**Table 1.** Participant demographics and clinical-neuropsychological scores.

|  | **Patients with schizophrenia** | **Healthy controls** | **P-value** |
| --- | --- | --- | --- |
|  | Mean±SD | Mean±SD |  |
| **N** | 19 | 19 |  |
| **Age [years]** | 41.1±12.17 | 36.84±11.5 | 0.27 |
| **Females/Males** | 7/12 | 6/13 | 0.73 |
| **FD [mm]** | 0.12±0.03 | 0.11±0.03 | 0.49 |
| **Injected radioactivity [MBq]** | 141.57±17.42 | 142.31±26.1 | 0.91 |
| **Illness duration [years]** | 13.47±9 | NA | NA |
| **Chlorpromazine equivalents [mg]** | 492.15±380.04 | NA | NA |
| **BACS [a.u.]** | 43.15±13.01 | 63.21±13.15 | <0.001 |
| **PANSS [a.u.]** |  |  |  |
| **Positive** | 11.36±3.53 | NA | NA |
| **Negative** | 14.31±6.36 | NA | NA |
| **General** | 26.73±7.42 | NA | NA |
| **Total** | 51.42±13.61 | NA | NA |

Age, FD, injected radioactivity, and BACS scores were compared with two-sample t-tests; gender via chi-squared test. Abbreviations: SD - standard deviation, N – number of participants, FD- framewise displacement, MBq – megabecquerel, BACS – Brief Assessment of Cognition in Schizophrenia, NA – not applicable, PANSS – Positive and Negative Syndrome Scale.

We investigated cortico-thalamic-iFC for SAL and ASM, respectively, with seed-based-iFC analysis restricted to the thalamus. Cortical seeds were derived from the 17-network cortical parcellation of Yeo and colleagues.^54^ Thalamic target regions-of-interest (ROI) were based on thalamic ROIs from our previous study of SAL-/ASM-hypo-/hyperconnectivity in an independent sample of patients with schizophrenia.^17^ Specifically, for SAL-iFC we masked the thalamus with the group-distinct cluster we previously reported to be hypoconnected with SAL in patients (mainly covering mediodorsal and ventral anterior nuclei), and for ASM-iFC we used the analogous cluster previously reported to be hyperconnected with ASM in patients (mainly covering ventral and posterior thalamic nuclei). Correlating the time-courses of these seeds and target-ROIs, we generated z-maps reflecting SAL-thalamic and ASM-thalamic-iFC (Figures 2B and 3B). Subsequently, we extracted voxel-wise iFC-values, averaged them across ROIs, and performed two-sample t-tests to compare the resulting ROI-wise iFC-values between groups. (For more details and description of control analyses (e.g. atlas-based thalamic ROIs) see Supplementary Methods.)

**Figure 2.**
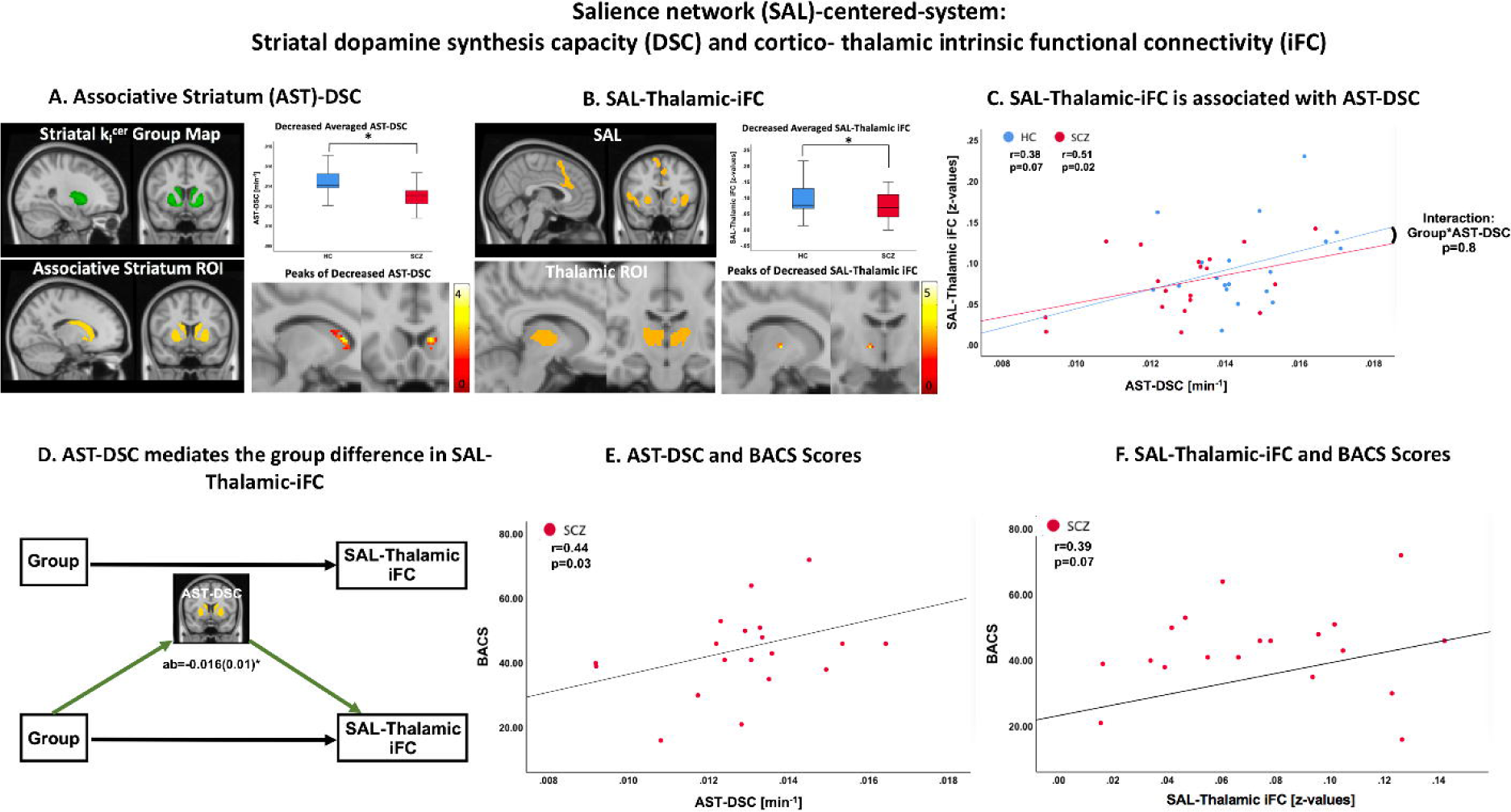
Salience network (SAL)-centered-system: Striatal dopamine synthesis capacity (DSC) and cortico-thalamic intrinsic functional connectivity (iFC) **A.** Associative striatum DSC. *Top left*: one-sample t-test parametric map of k_i_^cer^ (DSC) in associative striatum. *Top right*: box blots depicting significant group differences in associative-striatum-DSC, measured with a two-sample t-test on average DSC values (p=0.004). *Bottom left*: associative-striatum ROI from Oxford-GSK-Imanova atlas. *Bottom right*: two-sample t-test of voxel-wise group differences in associative-striatum-DSC; peak difference was found in left caudate (x=-12, y= 20, z=6; cluster-level corrected p_FWE_=0.05); color bar depicts range of t-values, with higher values in bright yellow. **B**. SAL-thalamic-iFC. *Top left*: template of prefrontal salience network.^54^ *Top right*: box blots depicting group differences in SAL-thalamic-iFC, measured with a two-sample t-test on average iFC values (p=0.04). *Bottom left*: thalamic ROI reflecting hypoconnectivity with SAL from our previous study.^17^ *Bottom right:* two-sample t-test of voxel-wise group differences in SAL-thalamic-iFC; peak difference was found in ventral anterior nucleus (x=12, y=-8, z=2; cluster-level corrected p_FWE_=0.05); color bar depicts range of t-values, with higher values in bright yellow. **C**. SAL-thalamic-iFC is associated with associative-striatum-DSC. Plot reflects partial correlation values (r) for associative-striatum-DSC and SAL-thalamic-iFC averaged values across patients with schizophrenia (red line and dots) and healthy controls (blue line and dots). Interaction effect between group and associative-striatum-DSC depicted with black curved line was not significant, p>0.05. **D**. Associative-striatum-DSC mediates the group difference in SAL-thalamic-iFC. *Top*: total effect – ‘path c’, reflecting the ‘simple’ association between the factor ‘group’ and SAL-thalamic iFC. *Bottom*: ‘path a’, reflecting effect of causal variable (group) on mediator (associative-striatum-DSC); ‘path b’, reflecting the effect of the mediator (associative-striatum-DSC) on the outcome variable (SAL-thalamic-iFC); ‘path ab’ – reflecting the indirect effect (mediation was significant), regression coefficient and standard error are shown; ‘path c’’, reflecting the direct effect of the causal variable (group) on the outcome variable (SAL-thalamic-iFC). **E**. Associative-striatum-DSC and BACS scores. Plot reflects partial correlation values (r) for averaged associative-striatum-DSC values and BACS scores across patients with schizophrenia. The correlation revealed a significant positive association (r=0.44, p=0.03). **F**. SAL-thalamic-iFC and BACS scores. Plot reflects partial correlation values (r) for averaged SAL-thalamic-iFC values and BACS scores across patients with schizophrenia. The correlation indicated a trend for positive association (r=0.39, p=0.07). *Abbreviations:* SAL – salience network, AST – associative-striatum, DSC – dopamine synthesis capacity, iFC – intrinsic functional connectivity, k_i_^cer^ – steady-state uptake rate constant of ^18^F-DOPA-PET relative to cerebellum concentration, HC – healthy controls, SCZ – patients with schizophrenia.

**Figure 3.**
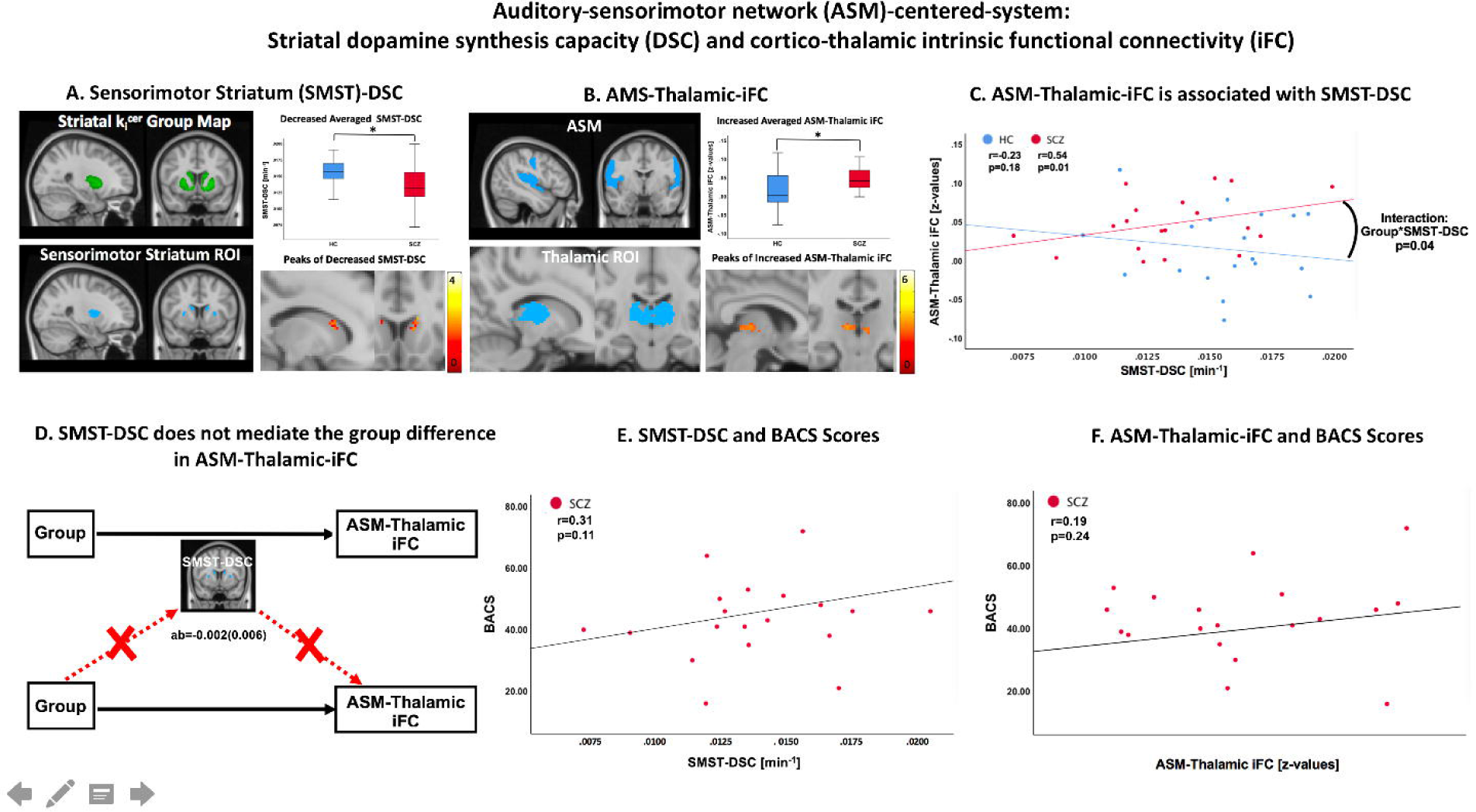
Auditory-sensorimotor network (ASM)-centered-system: Striatal dopamine synthesis capacity (DSC) and cortico-thalamic intrinsic functional connectivity (iFC) **A.** Sensorimotor striatum DSC. *Top left:* one-sample t-test parametric map of k_i_^cer^ (DSC) in sensorimotor-striatum. *Top right*: box blots depicting significant group differences in sensorimotor-striatum-DSC, measured with a two-sample t-test on average DSC values (p=0.02). *Bottom left:* sensorimotor-striatum ROI from Oxford-GSK-Imanova atlas. *Bottom right*: two-sample t-test of voxel-wise group differences in sensorimotor-striatum-DSC; peak difference was found in left putamen (x=-22, y=-4, z=6, uncorrected); color bar depicts range of t-values, with higher values in bright yellow. **B**. ASM-thalamic-iFC. *Top left*: template of auditory-sensorimotor network.^54^ *Top right*: box blots depicting group differences in ASM-thalamic-iFC, measured with a two-sample t-test on average iFC values (p=0.02). *Bottom left:* thalamic ROI reflecting hyperconnectivity with ASM from our previous study.^17^ *Bottom right*: two-sample t-test of voxel-wise group differences ASM-thalamic-iFC; peak difference was found in pulvinar (x=10, y=-24, z=4; cluster-level corrected p_FWE_=0.05); color bar depicts range of t-values, with higher values in bright yellow. **C**. ASM-thalamic-iFC is associated with sensorimotor-striatum-DSC. Plot reflects partial correlation values (r) for sensorimotor-striatum-DSC and ASM-thalamic-iFC averaged values across patients with schizophrenia (red line and dots) and healthy controls (blue line and dots). Interaction effect between group and sensorimotor-striatum-DSC depicted with black curved line was significant, p<0.05. **D**. Sensorimotor-striatum-DSC does not mediate the group difference in ASM-thalamic-iFC. *Top*: total effect – ‘path c’, reflecting the ‘simple’ association between the factor ‘group’ and ASM-thalamic iFC. *Bottom*: ‘path a’, reflecting effect of causal variable (group) on mediator (sensorimotor-striatum-DSC); ‘path b’, reflecting the effect of the mediator (sensorimotor-striatum-DSC) on the outcome variable (ASM-thalamic-iFC); ‘path ab’ – reflecting the indirect effect (mediation was *not* significant), regression coefficient and standard error are shown; ‘path c’’, reflecting the direct effect of the causal variable (group) on the outcome variable (ASM-thalamic-iFC). **E**. Sensorimotor-striatum-DSC and BACS scores. Plot reflects partial correlation values (r) for averaged sensorimotor-striatum-DSC values and BACS scores across patients with schizophrenia. The correlation was not significant (p=0.11). **F**. ASM-thalamic-iFC and BACS scores. Plot reflects partial correlation values (r) for averaged ASM-thalamic-iFC values and BACS scores across patients with schizophrenia. The correlation was not significant (p=0.24). *Abbreviations*: ASM – auditory-sensorimotor network, SMST – sensorimotor-striatum, DSC – dopamine synthesis capacity, iFC – intrinsic functional connectivity, k_i_^cer^ – steady-state uptake rate constant of ^18^F-DOPA-PET relative to cerebellum concentration, HC – healthy controls, SCZ – patients with schizophrenia.

### Statistical analyses for the relations between striatal-DSC, cortico-thalamic-iFC, and cognitive difficulties

Regarding our first hypothesis, we initially investigated relations between striatal-DSC and cortico-thalamic-iFC *within systems* (i.e., associative-striatum-DSC/SAL-thalamic-iFC and sensorimotor-striatum-DSC/ASM-thalamic-iFC, respectively) using partial correlation analyses with age, sex, head motion (FD), and chlorpromazine equivalents (CPZ) as covariates-of-no-interest. To investigate the relevance of DSC and iFC for patients’ impairments, i.e. their cognitive difficulties, we used the same approach to test whether DSC and iFC-measures of the SAL-centered and ASM-centered-systems, respectively, linked with BACS scores.

To foreshadow results, DSC and iFC-values were aberrant in patients and significantly correlated in both SAL- and ASM-centered-systems. To investigate whether these associations were distinct in the two groups, we (1) performed analogous partial correlation analyses also in healthy controls and (2) conducted interaction analyses (i.e., regression analyses testing for a significant effect of the interaction “group*striatal-subdivision-DSC” on the dependent variable cortico-thalamic-iFC).

Subsequently, to test whether aberrant striatal-DSC contributes to cortico-thalamic-dysconnectivity in schizophrenia, we used mediation analysis, i.e., we tested whether aberrant SAL-thalamic-iFC and ASM-thalamic-iFC was mediated by associative-striatum-DSC and sensorimotor-striatum-DSC, respectively.^55^ Mediation analysis investigates whether the effect of an independent variable ‘X’ (e.g., group) on a dependent variable ‘Y’ (e.g., SAL-thalamic-iFC) occurs via a third variable called mediator ‘M’ (e.g., associative-striatum-DSC).^56^ The relationships between these variables are quantified by regression analyses (we used age and gender as covariates-of-no-interest) and can be displayed as path diagrams (Figures 2D and 3D). The main outcome measure is the indirect effect ‘ab’, reflecting the product of X’s effect on M (‘a’) and M’s effect on Y (‘b’). The statistical significance of the indirect effect is determined via a nonparametric bootstrapping approach (we used 5,000 iterations) to obtain 95% confidence intervals.^55^ Together with corresponding hypotheses, mediation analysis provides both causal effects for longitudinal paradigms and conditional effects for cross-sectional approaches. Due to our cross-sectional approach, we used mediation analysis to test conditional relationships between schizophrenia, striatal-DSC, and cortico-thalamic-dysconnectivity; we applied the term ‘contributes to’ to depict significant mediation effects.

Finally, we tested our second hypothesis, whether SAL-centered-system changes contribute to ASM-centered-system changes. We used mediation analyses to investigate whether first, SAL-centered-system’s associative-striatum-DSC mediates the effect of schizophrenia on ASM-centered-system’s sensorimotor-striatum-DSC, and second, whether SAL-thalamic-iFC mediates the effect of schizophrenia on ASM-thalamic-iFC. Third, we conducted a three-way-interaction regression analysis to test for an influence of associative-striatum-DSC/SAL-thalamic-iFC on the group-distinct association between sensorimotor-striatum-DSC on ASM-thalamic-iFC (interaction “associative-striatum-DSC/SAL-thalamic-iFC*group*sensorimotor-striatum-DSC”, dependent variable: ASM-thalamic-iFC).

## Results

### SAL-centered-system: associative-striatum-DSC and SAL-thalamic-iFC

#### Decreased associative-striatum-DSC and SAL-thalamic-iFC in patients

Averaged associative-striatum-DSC values were significantly reduced (p=0.004) in patients (mean k_i_^cer^=0.0129±0.001 min^-1^) compared to healthy controls (mean k_i_^cer^=0.0144±0.001 min^-1^) (Figure 2A). Patients had reduced (p=0.04) SAL-mediodorsal/ventral-anterior-thalamus-iFC (mean z=0.07±0.04) compared to controls (mean z=0.10±0.05) (Figure 2B). In patients, no correlations were found between CPZ and DSC or iFC-values, respectively, indicating that decreased DSC and iFC-values were not confounded by current medication (see Supplementary Results for voxel-wise and further control analyses).

#### Decreased associative-striatum-DSC mediates SAL-thalamic-hypoconnectivity in patients

To study the topographic contribution of decreased associative-striatum-DSC to SAL-thalamic-hypoconnectivity, we first investigated the association between associative-striatum-DSC (k_i_^cer^) and SAL-thalamic-iFC (z-values) in patients and controls, respectively, via partial correlation analysis with age, sex, FD, and CPZ (only for patients) as covariates-of-no-interest. We found a significant positive correlation between associative-striatum-DSC and SAL-thalamic-iFC in patients (r=0.51, p=0.02) and at-trend-to-significant in controls (r=0.38, p=0.07) (Figure 2C). This association was not confounded by choice of thalamic ROIs (see Supplementary Results and Table S2). We found no interaction effect of “group*associative-striatum-DSC” on SAL-thalamic-iFC (p=0.8), indicating a continuum across individuals rather than a group-specific characteristic.

Critical to our first hypothesis, mediation analysis revealed a significant indirect effect of group on averaged SAL-thalamic-iFC via associative-striatum-DSC (ab=-0.016±0.01; 95%-confidence interval (CI) [−0.037 −0.0002]) (Figure 2D), indicating that reduced associative-striatum-DSC mediates reduced SAL-thalamic-iFC in patients. A control analysis demonstrated that SAL-thalamic-iFC did not mediate the effect of schizophrenia on associative-striatum-DSC (95%-CI [-0.009 0.0001]), indicating the specific contribution of associative-striatum-DSC to SAL-thalamic-iFC. In support of this specificity, no other significant mediations were found with several control analyses using alternative striatal subdivisions’ DSC as mediators. Additionally, current medication did not affect the contribution of associative-striatum-DSC to SAL-thalamic-iFC (see Supplementary Results).

#### Patients’ cognitive difficulties are linked with the SAL-centered-system

Due to previous findings,^17, 57, 58^ we expected that both aberrant associative-striatum-DSC and SAL-thalamic-iFC, respectively, are relevant for patients’ cognitive difficulties. To test this expectation, we used the partial correlation approach described above, with BACS scores reflecting patients’ cognitive difficulties. We found a positive correlation between BACS scores and associative-striatum-DSC (r=0.44, p=0.03) and at-trend-to-significance for SAL-thalamic-iFC (r=0.39, p=0.07), indicating the relevance of the SAL-centered-system for patients’ cognitive difficulties (Figure 2E and 2F; see Supplementary Results for control analyses).

### ASM-centered-system: sensorimotor-striatum-DSC and ASM-thalamic-iFC

#### Decreased sensorimotor-striatum-DSC and increased ASM-thalamic-iFC in patients

Averaged sensorimotor-striatum-DSC was significantly reduced (p=0.02) in patients (mean k_i_^cer^=0.0135±0.002 min^-1^) compared to controls (mean k_i_^cer^=0.0156±0.002 min^-1^) (Figure 3A). Patients had increased (p=0.02) ASM-ventral/posterior-thalamus-iFC (mean z=0.05±0.035) compared to controls (mean z=0.015±0.05) (Figure 3B). In patients, no correlations were found between CPZ, DSC, and iFC-values, respectively, indicating that decreased DSC and increased iFC-values were not confounded by current medication (see Supplementary Results for voxel-wise and control analyses).

#### Sensorimotor-striatum-DSC is positively associated with ASM-thalamic-iFC in patients, but decreased sensorimotor-striatum-DSC does not mediate ASM-thalamic-hyperconnectivity

Next, we tested for the topographic contribution of reduced sensorimotor-striatum-DSC to ASM-thalamic-hyperconnectivity. We first investigated the association between sensorimotor-striatum-DSC (k_i_^cer^) and ASM-thalamic-iFC (z-values) using the partial correlation approach. A significant positive correlation between sensorimotor-striatum-DSC and ASM-thalamic-iFC was found in patients (r=0.54, p=0.01) but not in controls (r=-0.23, p=0.18) (Figure 3C). This association was not confounded by the choice of thalamic ROIs (see Supplementary Results and Table S2). Furthermore, we found a significant interaction between group and sensorimotor-striatum-DSC (p=0.04), indicating that – for ASM-centered-system – sensorimotor-striatum-DSC and ASM-thalamic-iFC are group-distinctively related.

Next, we employed mediation analysis to test whether the effect of schizophrenia on ASM-thalamic-iFC was mediated by sensorimotor-striatum-DSC. The indirect effect was not significant (95%-CI [-0.02 0.01]) (Figure 3D). Furthermore, ASM-thalamic-iFC did not mediate the effect of schizophrenia on sensorimotor-striatum-DSC either (95%-CI [-0.017 0.01]. These results indicate that although sensorimotor-striatum-DSC is associated with ASM-thalamic-iFC, the effect of the schizophrenia on ASM-thalamic-hyperconnectivity is not mediated by sensorimotor-striatum-DSC. Several control analyses, in which alternative striatal subdivisions’ DSC were used as mediator variables, showed that striatal-DSC does not contribute to ASM-thalamic-iFC directly (see Supplementary Results). As ASM-thalamic-iFC did also not mediate the group effect on sensorimotor-striatum-DSC, these results suggest a potential third factor inducing the patient-specific association between sensorimotor-striatum-DSC and ASM-thalamic-iFC (see below).

#### Patients’ cognitive difficulties do not link with the ASM-centered-system

Based on previous findings,^17^ we did not expect an association between ASM-centered system and patients’ cognitive difficulties. We investigated the corresponding relationship, using the same partial correlation approach as described for SAL-centered-system. We did not find any association between the ASM-centered-system and cognitive difficulties in patients with partial correlation analyses. Specifically, no significant correlation was found between patients’ BACS scores and sensorimotor-striatum-DSC (r=0.31, p=0.11) or ASM-thalamic-iFC (r=0.19, p=0.24) (Figure 3E and 3F). Control analyses demonstrated that choice of thalamic ROIs did not confound these results (see Supplementary Results).

### Relationship between SAL- and ASM-centered-systems

According to our second hypothesis, we expected SAL-centered-system changes to contribute to changes in the ASM-centered-system.

#### Associative-striatum-DSC mediates the group difference in sensorimotor-striatum-DSC

Mediation analysis revealed a significant indirect effect of group on sensorimotor-striatum-DSC via associative-striatum-DSC (ab=-0.002±0.001; 95%-CI [−0.005 −0.0005]) (Figure 4 and S3A), indicating that reduced associative-striatum-DSC mediates the effect of schizophrenia on sensorimotor-striatum-DSC. A control-analysis testing for the opposite effect, however, revealed that sensorimotor-striatum-DSC also mediates the group effect on associative-striatum-DSC, indicating both anterior-to-posterior and posterior-to-anterior effects (see Supplementary Results).

**Figure 4.**
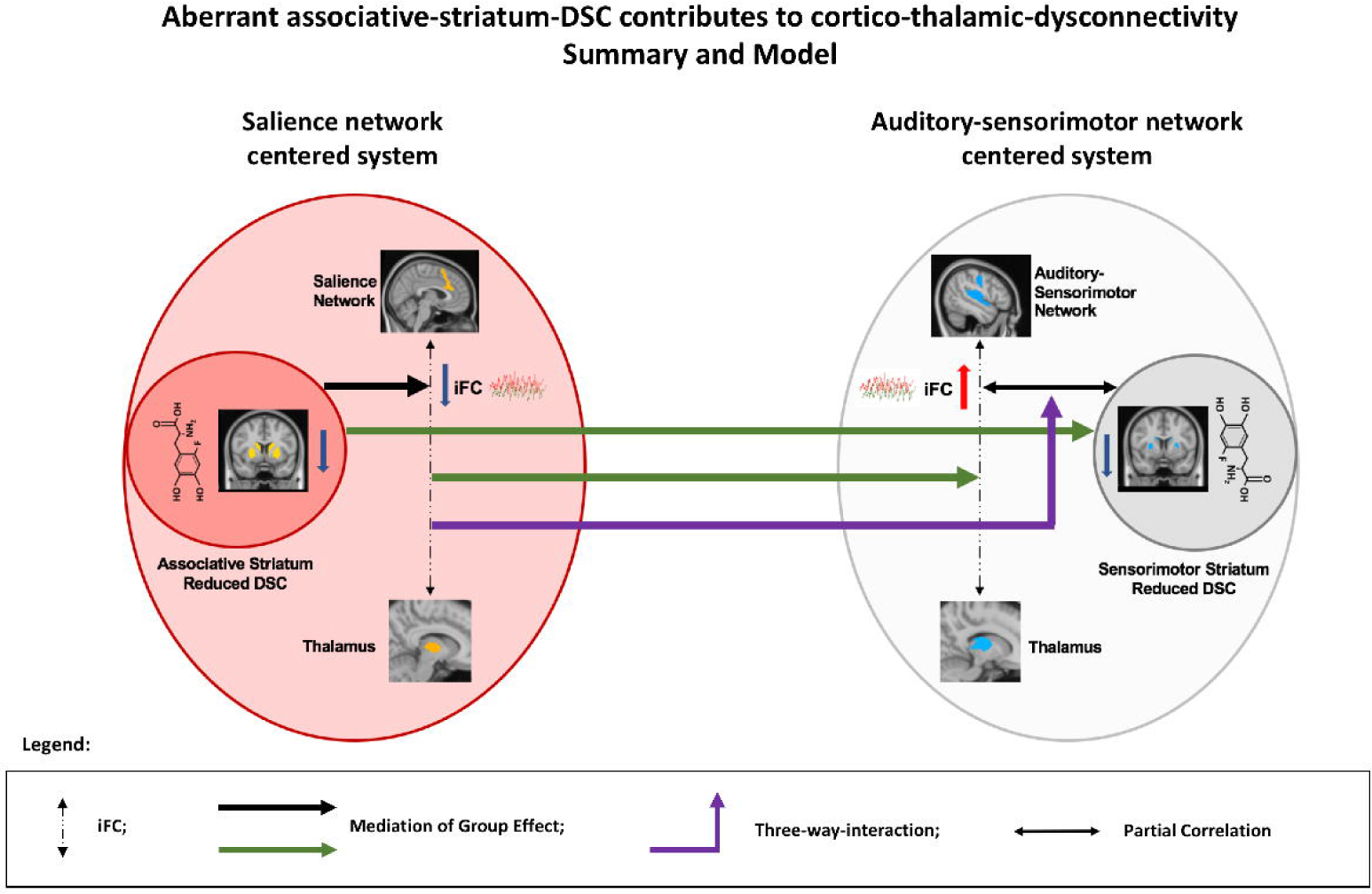
Aberrant associative-striatum-DSC contributes to cortico-thalamic-dysconnectivity. Summary and Model. Arrows (see legend below) represent a summary of results regarding the observed relationships between SAL- and ASM-centered-systems. The systems’ colors represent the model derived from these results: associative-striatum dopamine has a primary role for cortico-thalamic-dysconnectivity (for details see discussion). *Left*: The SAL-centered-system is depicted (light-red circle) with its two components, associative-striatum-DSC (dark-red circle) and SAL-thalamic-iFC. Both associative-striatum-DSC and SAL-thalamic-iFC were reduced in patients as shown by blue arrows pointing down. Reduced associative-striatum-DSC mediated SAL-thalamic-hypoconnectivity in patients, depicted by the black arrow from associative-striatum-DSC to SAL-thalamic-iFC. *Right*: The ASM-centered-system is depicted with its two components, sensorimotor-striatum-DSC (grey circle) and ASM-thalamic-iFC. Sensorimotor-striatum-DSC was reduced in patients as shown by blue arrow pointing down and ASM-thalamic-iFC was increased as shown by red arrow pointing up. In patients, the association between reduced sensorimotor-striatum-DSC and increased ASM-thalamic-iFC was significant, depicted by double black arrow. *Central:* relations between SAL- and ASM-*centered-systems*. Associative-striatum-DSC mediated the group difference in sensorimotor-striatum-DSC, shown as green arrow from left to right; SAL-thalamic-iFC mediated the group difference in ASM-thalamic-iFC, also shown as green arrow from left to right; and the group-distinct association between reduced sensorimotor-striatum-DSC and increased ASM-thalamic-iFC was moderated by reduced SAL-thalamic-iFC, shown as purple arrow from left to right. *Abbreviations*: DSC – dopamine synthesis capacity, iFC – intrinsic functional connectivity.

#### SAL-Thalamic-iFC mediates the group difference in ASM-Thalamic-iFC

Furthermore, mediation analysis revealed a significant indirect effect of group (patients and healthy controls) on ASM-thalamic-iFC via SAL-thalamic-iFC (ab=-0.018±0.001; 95%-CI [−0.038 − 0.0003]) (Figure 4 and S3B), demonstrating that reduced SAL-thalamic-iFC mediates increased ASM-thalamic-iFC in patients. The opposite did not hold, as ASM-thalamic-iFC does not mediate the group effect on SAL-thalamic-iFC, indicating a specific anterior-to-posterior effect (see Supplementary Results).

#### SAL-Thalamic-iFC significantly moderates the group-distinct association between sensorimotor-striatum-DSC and ASM-Thalamic-iFC

Finally, we tested whether associative-striatum-DSC and SAL-thalamic-iFC, respectively, moderate the group difference in the association between sensorimotor-striatum-DSC and ASM-thalamic-iFC. A three-way-interaction demonstrated that only SAL-thalamic-iFC (p=0.02) but not associative-striatum-DSC (p=0.15) moderated the interaction between group and sensorimotor-striatum-DSC on ASM-thalamic-iFC (Figure 4 and S3C), indicating that the patient-specific association between sensorimotor-striatum-DSC and ASM-thalamic-iFC is moderated by SAL-thalamic-iFC.

## Discussion

Using simultaneous ^18^F-DOPA-PET and rs-fMRI in patients with schizophrenia during remission of positive symptoms, we provide first-time evidence that aberrant striatal dopamine in schizophrenia links with cortico-thalamic-dysconnectivity. Particularly, in the salience-network-centered-system, aberrant dopamine synthesis capacity in the associative-striatum mediated patients’ decreased SAL-mediodorsal-thalamus intrinsic functional connectivity, while in the auditory-sensorimotor-network-centered-system, such a mediation was absent. Furthermore, alterations in the SAL-centered-system (regarding both striatal-DSC and cortico-thalamic-iFC) mediated corresponding changes in the ASM-centered-system. The mediating role of aberrant associative-striatum-DSC for SAL-thalamic-dysconnectivity together with the anterior-to-posterior gradient of contribution from the SAL-to the ASM-centered-system suggest a central role of associative-striatum dopamine dysfunction in the pathophysiology of schizophrenia.

### Decreased associative-striatum-DSC, but not decreased sensorimotor-striatum-DSC, mediates cortico-thalamic-dysconnectivity

We expected aberrant striatal-DSC to topographically mediate cortico-thalamic-dysconnectivity. First, in the SAL-centered-system, we found both associative-striatum-DSC and SAL-thalamic-iFC reduced in patients (Figure 2A, 2B), while in the ASM-centered-system, patients’ sensorimotor-striatum-DSC was reduced but ASM-thalamic-iFC increased (Figure 3A, 3B). These results were basic for our study and replicated several previous reports for both aberrant DSC^8, 25^ and iFC.^13, 17, 30^

In the SAL-centered-system, decreased associative-striatum-DSC was positively associated with SAL-thalamic-iFC in patients and at-trend-to-significance in healthy controls (Figure 2C). An interaction analysis demonstrated that this association was not schizophrenia-specific (Figure 2C), indicating rather a continuum across individuals. This finding is supported by a recent study, which reported a specific association between within-SAL-iFC and associative-striatum-DSC in healthy controls.^38^ Critically and in line with our expectation, aberrant associative-striatum-DSC mediated the effect of schizophrenia on SAL-thalamic-iFC (Figure 2D). This mediation was specific for associative-striatum-DSC, as no other striatal subdivision DSC had a mediating effect on SAL-thalamic-iFC, nor did SAL-thalamic-iFC mediate the effect of schizophrenia on associative-striatum-DSC. These results were not confounded by current medication, head motion, or choice of thalamic/striatal ROIs, as we controlled for these factors. Therefore, decreased associative-striatum-DSC contributes to SAL-thalamic-hypoconnectivity in schizophrenia, and might actually drive SAL-thalamic-dysconnectivity. However, to confirm this last suggestion, longitudinal studies are needed.

In the ASM-centered-system, in turn, decreased sensorimotor-striatum-DSC was positively associated with ASM-thalamic-iFC in patients but not in healthy controls (Figure 3C). An interaction analysis demonstrated that this association was significantly distinct for both groups, and therefore schizophrenia-specific (Figure 3C). The missing association in healthy controls is in line with the above-mentioned study, which did not find any association between striatal-DSC and sensorimotor-network-iFC.^38^ Contrary to our hypothesis, we found that sensorimotor-striatum-DSC did not mediate the effect of schizophrenia on ASM-thalamic-iFC (Figure 3D). Neither did ASM-thalamic-iFC mediate group effects on sensorimotor-striatum-DSC, suggesting a further third factor – presumably from the SAL-centered-system – as potentially relevant for the schizophrenia-specific association between sensorimotor-striatum-DSC and ASM-thalamic-iFC (see below).

Our main finding – decreased associative-striatum-DSC contributing to SAL-thalamic-hypoconnectivity in schizophrenia - is congruent with general evidence that the dopaminergic modulated basal ganglia influence cortico-thalamic-connectivity^18, 20, 59^ through parallel, topographically organized cortico-striato-pallido-thalamo-cortical circuits (CSPTC).^42, 60, 61^ Particularly, associative cortices including frontal/SAL regions (i.e., opercular-insula, dorsal anterior cingulate) form, together with associative-striatum and mediodorsal nucleus, parts of the *associative* CSPTC-circuit.^34, 62^ The circuit is modulated by dopaminergic projections from substantia nigra pars compacta, which projects topographically in a rostral-to-caudal gradient to cortical and subcortical areas, including dorsal striatum and mediodorsal nucleus.^63-66^ Therefore, associative-striatum-DSC changes are relevant for associative CSPTC-circuit aberrances - partly represented by SAL-thalamic-dysconnectivity - in schizophrenia, and aberrant nigro-striatal dopaminergic projections might have a primary role in the pathophysiology of the associative CSPTC-circuit in schizophrenia.

### SAL-centered-system changes contribute to ASM-centered-system changes in schizophrenia

Concerning our second hypothesis, we identified three pathways through which SAL-centered-system changes contribute to ASM-centered-system changes. In particular, the first two of these pathways demonstrate a direct anterior-to-posterior mediation effect. First, decreased associative-striatum-DSC mediated the effect of schizophrenia on sensorimotor-striatum-DSC (Figure S3A), indicating that aberrant dopamine in the associative part contributes to dopamine changes in the sensorimotor part of the striatum, being in line with previous findings^40, 46^. This effect was not specific, as sensorimotor-striatum-DSC also mediated the effect of schizophrenia on associative-striatum-DSC, suggesting that dorsolateral striatum is also relevant for dorsomedial striatum dopamine function. This last finding is in line with recent reports regarding striatal dorsolateral-to-dorsomedial influences (see Supplemental Discussion).^46^ Second, decreased SAL-thalamic-iFC mediated the effect of schizophrenia on increased ASM-thalamic-iFC (Figure S3B). This effect was specific, as ASM-thalamic-iFC did not mediate the group difference in SAL-thalamic-iFC. Results are supported by physiological evidence that (i) in the striatum ventral/dorsomedial areas (including associative-striatum) affect more dorsolateral areas (including sensorimotor-striatum) by a variety of dopamine-dependent mechanisms, enabling topographical crosstalk between different CSPTC-circuits^40, 43, 46^ – i.e., ascending spiral of striato-nigro-striatal projections^43, 47^ - and (ii) direct cortico-cortical and indirect trans-thalamic cortico-thalamic-cortical influences, possibly underpinning anterior-to-posterior crosstalk between CSPTC-circuits.^20, 59, 67^

The third pathway demonstrated an indirect anterior-to-posterior effect. Specifically, a three-way-interaction revealed that reduced SAL-thalamic-iFC, but not associative-striatum-DSC, moderated the interaction between group and sensorimotor-striatum-DSC on ASM-thalamic-iFC (Figure S3C), indicating an additional anterior-to-posterior effect through altered SAL-centered-system iFC. Notably however, aberrant SAL-thalamic-iFC was in turn mediated by decreased associative-striatum-DSC, suggesting a primary role for associative-striatum dopamine dysfunction in these pathophysiological relationships between SAL- and ASM-centered-systems.

### Model: Aberrant associative-striatum-DSC contributes to cortico-thalamic-dysconnectivity

In summary, our findings suggest an integrative model in which the SAL-centered-system exerts a three-pathway influence over the ASM-centered-system: first, reduced associative-striatum-DSC mediated reduced sensorimotor-striatum-DSC in patients (Figure 4, upper green arrow). Second, decreased SAL-thalamic-iFC mediated increased ASM-thalamic-iFC in patients (Figure 4, lower green arrow). Third, SAL-thalamic-iFC moderated the group-distinct association between sensorimotor-striatum-DSC and ASM-thalamic-iFC (Figure 4, purple arrow). Within the SAL-centered-system, on the other hand, associative-striatum-DSC mediated SAL-thalamic-hypoconnectivity in patients (Figure 4, black arrow from associative-striatum-DSC to SAL-thalamic-iFC). This suggests altered associative-striatum-DSC as the main driver of pathophysiological changes in both SAL- and ASM-centered-systems. The associative-striatum is an extraordinary information processing hub in the brain that receives and integrates massive and highly heterogeneous cortical inputs^68-70^ and facilitates crosstalk between CSPTC-circuits.^40^ Correspondingly, aberrant associative-striatum function, driven by aberrant associative-striatum-dopamine, might have widespread pathophysiological effects across distinct systems such as within SAL- and into ASM-centered system. These broad changes likely underpin widespread functional impairments.^71^ Indeed, we found a specific association of both associative-striatum-DSC and SAL-thalamic-iFC with patients’ cognitive difficulties (see Supplementary Discussion for more details).

### Strengths and limitations

#### Strengths

(i) The current study is the first to investigate relations between striatal-DSC and cortico-thalamic-iFC with simultaneous ^18^F-DOPA-PET and rs-fMRI, avoiding temporal confounds of serial assessment. (ii) We recruited a well-defined, homogenous sample of patients, avoiding confounds of symptomatic variance. (iii) Our results lead directly to an integrating model, supporting current and previous results and specifying testable hypotheses. Particularly, this model integrates two largely independent lines of findings in schizophrenia: striatal dopamine dysfunction and cortico-thalamic dysconnectivity.

#### Limitations

(i) Patients were medicated with anti-psychotic drugs, potentially inducing confounding effects (Table S1). Although we did not find such (current) medication confounds with several control analyses, results should be nevertheless evaluated carefully, for example we were not able to control for effects of past medication. (ii) The interpretation of k_i_^cer^ as index of DSC needs caution, as tyrosine hydroxylase is the rate-limiting step in DSC and not aromatic acid decarboxylase (for further limitations regarding the DSC measure such as premedication see Supplementary Discussion).^6, 72^ (iii) Striatal-DSC and blood oxygenation-based cortico-thalamic-iFC, are indirect measures of brain activity, averaged over several minutes and reflecting mixed signals. Therefore, we are only able to examine relative coarse relations and we might have missed fast occurring effects (e.g. of phasic dopaminergic transmission).

## Conclusion

Cortico-thalamic-dysconnectivity links with aberrant striatal dopamine in schizophrenia.

## Acknowledgements

We thank Sylvia Schachoff and Anna Winter for their technical assistance during PET/MRI measurements. Particularly, we thank all subjects for participating in the study.

## Funding

This work has been supported by the European Union 7^th^ Framework Programme, TRIMAGE, a dedicated trimodality (PET/MR/EEG) imaging tool for schizophrenia (Grant no. 602621).

## Competing interests

The authors report no conflict of interest relating to this work.

S.L. has received honoraria for consulting or lectures from LB Pharma, Lundbeck, Otsuka, TEVA, LTS Lohmann, Geodon Richter, Recordati, Boehringer Ingelheim, Sandoz, Janssen, Lilly, SanofiAventis, Servier, and Sunovion.

