## Supplementary material for "Cortico-thalamic dysconnectivity links with aberrant striatal dopamine in schizophrenia *A simultaneous ^18^F-DOPA-PET/resting-state fMRI study*"

By Avram, Brandl et al.

### **Supplementary Methods**

#### Participants

23 patients meeting DSM-IV criteria for schizophrenia (age range: 23-65 years; mean: 43.04±11.90 years) and 24 healthy subjects comparable for age and sex (age range: 25-62 years; mean: 38.54±11.63 years) were recruited from the Department of Psychiatry, Klinikum rechts der Isar of the Technical University of Munich, Germany, and from the Munich area by advertising, respectively. Due to excessive head motion during the fMRI resting-state sequence, several subjects were excluded from both groups, with 19 subjects remaining in each group for further analyses. Clinical and neuropsychological assessment was carried out by the same trained rater (C.L.) for all participants. Psychiatric diagnosis was supported by the Structured Clinical Interview for DSM-IV.^1^ The positive and negative syndrome scale (PANSS) was used to assess symptom severity.^2^ Cognitive abilities were evaluated with the Brief Assessment of Cognition in Schizophrenia (BACS), a test battery developed to assess cognitive performance in schizophrenia.^3^ All patients had established schizophrenia (at least 2 previous psychotic episodes) and were in symptomatic remission of positive symptoms according to the criteria of Andreasen and colleagues^~~4~~^ at the time of the scan – i.e., patients' scores on PANSS items ’delusions’ (P1), ‘conceptual disorganization’ (P2), 'hallucinatory behavior' (P3), 'mannerisms/posturing' (G5), and 'unusual though content' (G9) were ≤3.^4, 5^ However, no remission criteria had to be fulfilled for negative and general symptoms. Before scanning, antipsychotic medication was kept stable for a minimum of 2 weeks due to temporal dynamics of elimination and effects on DSC (see Table S1 for detailed antipsychotic and adjunctive psychotropic medication). Patients with substance abuse (besides nicotine) were excluded from the study. Healthy controls did not have a history of psychiatric illness, substance abuse, or first-degree relatives with schizophrenia. Participants gave their written informed consent after receiving a complete description of the study. The study was approved by the Ethics Review Board of the Technical University of Munich, and approval to administer radiotracers was obtained from the Administration of Radioactive Substances (Bundesamt für Strahlenschutz), Germany.

#### ^18^F-DOPA-PET data acquisition, preprocessing, and k_i_^cer^ as outcome measure for dopamine synthesis capacity (DSC)

##### *^18^F-DOPA-PET data reconstruction*

^18^F-DOPA-PET and MRI data were acquired simultaneously with a hybrid whole-body mMR Biograph PET/MRI scanner (Siemens-Healthineers, Erlangen, Germany), using a vendor-supplied 12-channel phase-array head coil. ^18^F-DOPA-PET measures the influx constant k_i_^cer^, a quantitative measure reflecting DSC. Participants were instructed not to smoke, drink coffee or alcohol for 12 hours before scanning,^6^ and received about 150 MBq of ^18^F-DOPA (mean: 140.25±21.15 MBq). PET acquisition lasted 70min. Ordered subset expectation maximization (OSEM) (21 subsets, 3 iterations) was used to reconstruct PET data with a voxel size of 1.7×1.7×2mm, and a 3mm Gaussian post-reconstruction filter, corrected for attenuation^7^ and scatter based on anatomical MRI information. The reconstruction was performed using the Siemens e7 off-line PET reconstruction toolkit, which included attenuation, scatter, randoms, and decay corrections, and normalization. PET data were framed into 30 dynamic frames (1×30s, 10×15s, 3×20s, 2×60s, 2×120s, 12×300s). We used the cerebellum as reference region in a voxel-wise manner with Gjedde–Patlak linear graphical analysis.^8^ PET frames acquired between 20-60 minutes were used for linear fit, which resulted in whole-brain voxel-wise k_i_^cer^ maps*.* We computed mean k_i_^cer^ by averaging over all voxels in the striatum and striatal subdivisions (limbic, associative, and sensorimotor – see below).

PET data were analyzed together with the T1-weighted MRI data (see below) using Statistical Parametric Mapping, version 12 (SPM12: https://www.fil.ion.ucl.ac.uk/spm/). Briefly, PET data were corrected for motion by realigning all PET frames to the last frame. Each individual T1-image was co-registered to the last PET frame, followed by spatial normalization into MNI space. Subsequent to the normalization, the inverse transformation matrix was applied to all regions-of-interest (ROIs) to transform them into each individual’s PET space.

##### *Regions-of-interest and masks used for the generation of whole-brain voxel-wise k_i_^cer^ maps*

We used FSL to create an anatomical mask of the cerebellum (excluding vermis) from the probabilistic cerebellar atlas,^9^ and extracted its time activity curve. Mean k_i_^cer^ values of the striatum and its subdivisions were extracted (by averaging over all voxels within each region-of-interest) from the k_i_^cer^ maps using functionally defined masks from the Oxford-GSK-Imanova connectivity atlas (limbic, associative, and sensorimotor subdivisions) and their sum for whole striatum. We then tested for between-group effects regarding average DSC of striatal subdivisions with two-sample t-tests by using SPSS Statistics for Macintosh, Version 25.0. (IBM Corp. 2017).

*Control analyses with voxel-wise approach*

To control for this regional approach, and to locate peak differences, we normalized the k_i_^cer^ maps into MNI space with SPM12, and performed voxel-wise statistical analyses by entering the normalized k_i_^cer^ maps into one- and two-sample voxel-wise t-tests with SPM12 (cluster-level p_FWE_=0.05), using age and gender as covariates of no interest. For one-sample t-maps for each group see Supplement (Figure S1).

#### MRI data acquisition, preprocessing, and correlation coefficients as outcome measure for intrinsic functional connectivity (iFC)

##### *Control analyses with voxel-wise approach*

To control for the ROI-based approach and to locate peaks of group difference, we performed voxel-wise one- and two-sample tests for SAL-thalamic and ASM-thalamic-iFC maps with SPM12 (https://www.fil.ion.ucl.ac.uk/spm/), with cluster-level p_FWE_=0.05 thresholding for voxel-wise p=0.001 and age and sex as covariates of no interest. For one-sample t-maps for each group see Supplement (Figure S1).

##### *Control analyses with anatomically defined thalamic ROIs*

To control for possible effects of the thalamic ROIs on the associations between iFC and DSC measures, as well as between them and cognitive scores, we extracted iFC-values from thalamic nuclei defined anatomically with WFU Pickatlas (http://fmri.wfubmc.edu/software/pickatlas) instead of the hypo-/hyperconnectivity clusters based on our previous study,^10^ in the following manner: for SAL-iFC we used the mediodorsal nucleus and ventral anterior nucleus, since both nuclei have been shown to have iFC with SAL^11^ (for orientation see both thalamic ROIs in Figure S2, Panel A); for ASM-iFC we used the pulvinar and ventrolateral nucleus, since both nuclei have been shown to have iFC with ASM^11^ (for orientation see both thalamic ROIs in Figure S2, Panel B). We then extracted and averaged for each individual, on the one hand, iFC-values from thalamic voxels of the aforementioned thalamic nuclei (mediodorsal and ventral anterior for SAL-thalamic-iFC, pulvinar and ventrolateral nucleus for ASM-thalamic-iFC), and on the other hand, k_i_^cer^-values from corresponding k_i_^cer^ maps of the striatum and striatal subdivisions, respectively. We then entered these averaged values in partial correlation analyses using age, sex, and FD as covariates-of-no-interest for both groups, and also chlorpromazine equivalents (CPZ) for the patient group.

#### Statistical analyses for the link between iFC, DSC, and cognitive difficulties

##### *Control analyses regarding the directional specificity of the influence of DSC on iFC*

In order to test whether striatal subdivisions’ DSC contributes to cortico-thalamic-dysconnectivity and not the other way around, we repeated the mediation analyses by using striatal subdivisions’ DSC as outcome variable and cortico-thalamic iFC as mediator variable. Specifically, we tested whether, on the one hand, SAL-thalamic-iFC mediates the effect of schizophrenia on associative-striatum-DSC, and on the other hand, whether ASM-thalamic-iFC mediates the effect of schizophrenia on sensorimotor-striatum-DSC.

##### *Specificity and confirmatory analyses: striatal subdivisions, group, and thalamic ROIs specificity*

To control and explore regional specificity of the link between iFC and striatal subdivisions’ DSC, we repeated the partial correlation analyses across patients by replacing the subdivision of interest (e.g. associative-striatum-DSC) with an alternative subdivision (e.g. sensorimotor-striatum-DSC) or with whole-striatum-DSC. To control for potential confounds elicited by thalamic ROI definition, we repeated the partial correlation analyses for iFC values based on alternative thalamic ROI values. Specifically, in line with the previously mentioned control analyses, we tested for associations between: (1) associative-striatum-DSC and SAL-mediodorsal-iFC and also for associative-striatum-DSC and SAL-ventral-anterior-iFC in both patients and healthy controls; (2) sensorimotor-striatum-DSC and ASM-pulvinar-iFC and also for sensorimotor-striatum-DSC and ASM-ventrolateral-iFC in both patients and healthy controls; (3) SAL-mediodorsal-iFC and BACS scores as well as SAL-ventral-anterior-iFC and BACS scores, in both patients and healthy controls.

### **Supplementary Results**

#### SAL-centered-system: associative-striatum-DSC and SAL-thalamic-iFC

##### *Voxel-wise and control analyses (movement and current medication) for both associative-striatum-DSC and SAL-thalamic-iFC*

After establishing that averaged associative-striatum-DSC was reduced in patients, we controlled for this regional approach with a voxel-wise two-sample t-test within the associative-striatum-ROI, which revealed the peak of group difference in the left caudate (x=-12, y= 20, z=6; cluster-level corrected p_FWE_=0.05), indicating consistent DSC reductions in the associative-striatum. Furthermore, there was no significant association between averaged associative-striatum-DSC and CPZ (r=0.05, p=0.83) or FD (r=-0.22, p=0.536) in patients, suggesting that associative-striatum-DSC reductions were not confounded by current medication or head motion.

We found that averaged SAL-thalamic-iFC was reduced in patients. This result was supported by a voxel-wise two-sample t-test, which revealed peaking SAL-thalamic-hypoconnectivity in the ventral anterior nucleus (x=12, y=-8, z=2; cluster-level corrected p_FWE_=0.05 for voxel-wise p=0.001). Furthermore, no significant associations between SAL-thalamic-iFC and CPZ (r=0.35, p=0.14) or FD (r=0.13, p=0.57), were found in the patient group, as shown by Pearson’s correlation analyses, suggesting that SAL-thalamic-iFC results were not confounded by current medication or head motion

##### *Further control analyses for whole striatum DSC and limbic striatum DSC*

As confirmatory analyses, we also investigated averaged DSC in whole striatum and limbic-striatum subdivision. Whole striatum DSC was reduced (p=0.01) in patients (mean 0.0130±0.001 min^-1^) compared to healthy controls (mean 0.0142±0.001 min^-1^). No differences were found for limbic-striatum-DSC (p=0.21) between patients (mean 0.0123±0.001 min^-1^) and healthy controls (mean 0.0129±0.001 min^-1^). Both results are in line with our previous findings in a larger sample.^12^ Next, we tested for associations between whole striatum DSC and limbic-striatum-DSC and CPZ and FD, respectively. There was no significant association between averaged whole striatum DSC and CPZ (r=0.37, p=0.12) or FD (r=-0.18, p=0.45) in patients, suggesting that striatal-DSC reductions are not confounded by current medication or head motion. There was a trend association for limbic-striatum-DSC and CPZ (r=0.44, p=0.054), which, however, was not used in any analysis. No association was found between limbic-striatum-DSC and FD (r=-0.14, p=0.56).

##### *Decreased associative-striatum-DSC mediates decreased SAL-thalamic-iFC in patients*

##### *Associations between associative-striatum-DSC and SAL-thalamic-iFC using alternative thalamic ROIs*

To control for the effect of the thalamic ROI, we tested for associations between SAL-thalamic-iFC and associative-striatum-DSC in patients by using anatomically defined thalamic nuclei instead of the hypoconnectivity cluster from our previous study.^10^ We found very similar results as before, namely SAL-mediodorsal-nucleus-iFC correlated significantly with associative-striatum-DSC (r=0.53, p=0.02) and SAL-ventral-anterior-nucleus-iFC at-trend-for-significance with associative-striatum-DSC (r=0.41, p=0.06) (Table S2). We conclude that our finding regarding the association between associative-striatum-DSC and SAL-thalamic-iFC was not confounded by the choice of thalamic ROI.

Subsequently, to control for the specificity of associative-striatum-DSC to being relevant for SAL-thalamic-iFC, we tested whether SAL-thalamic-iFC also correlates with other striatal subdivisions’ DSC or whole striatum DSC. We found a significant correlation between sensorimotor-striatum-DSC and SAL-thalamic-iFC in patients (r=0.48, p=0.03), but not limbic-striatum-DSC (r=0.30, p=0.13) or whole striatal-DSC (r=0.13, p=0.31). These findings suggest SAL-thalamic-iFC to be associated with the dorsal striatum (associative- and sensorimotor striatum), however, the highest correlation was found for associative-striatum-DSC.

##### *Mediation analyses for additional striatal subdivisions’ DSC*

After establishing that associative-striatum-DSC mediates the effect of schizophrenia on SAL-thalamic-iFC, we tested whether the other striatal subdivisions have a similar effect. For whole striatal-DSC, mediation analysis revealed that the indirect effect was not significant (the bootstrapped 95% confidence interval [-0.03 0.004] included zero). Similar, for limbic-striatum-DSC, mediation analysis revealed that the indirect effect was not significant (95% confidence interval [-0.017 0.005] included zero). Finally, for sensorimotor-striatum-DSC, mediation analysis revealed that the indirect effect was also not significant (95% confidence interval [-0.03 0.001] included zero). We conclude that specifically associative-striatum-DSC mediates the group difference in SAL-thalamic-iFC.

##### *Control for effects of current medication on the mediation*

To test whether the mediation was confounded by effects of current medication, we repeated the mediation analysis – to test whether the effect of schizophrenia on SAL-thalamic-iFC was mediated by associative-striatum-DSC - but added CPZ to the covariates-of-no-interest. The indirect effect was still significant (ab=-0.017±0.01; the bootstrapped 95% confidence interval [-0.04 -0.0002] did not include zero), suggesting that current medication does not affect the contribution of associative-striatum-DSC to SAL-thalamic-iFC.

##### *Patients’ cognitive difficulties link with the SAL-centered-system*

##### *Control for the influence of alternative thalamic ROIs on the association between SAL-thalamic-iFC and BACS scores*

To test whether anatomically defined thalamic ROIs could affect the association between SAL-thalamic-iFC and BACS scores in patients, we repeated the partial correlation analysis using the mediodorsal and ventral anterior nucleus, respectively, instead of the hypoconnectivity cluster from our previous study.^10^ We found SAL-mediodorsal-thalamus-iFC to be correlated with BACS scores (r=0.44, p=0.05) but not SAL-ventral-anterior-thalamus-iFC (r=0.27, p=0.16). These findings indicate that the mediodorsal nucleus – which is also included in the hypoconnectivity cluster (Figure S2) – is likely driving the association between SAL-thalamic-iFC and BACS scores in patients. See Table S2 for correlations in healthy controls.

#### ASM-centered-system: sensorimotor-striatum-DSC and ASM-thalamic-iFC

##### *Voxel-wise and control analyses for sensorimotor-striatum-DSC and ASM-thalamic-iFC*

After establishing that averaged sensorimotor-striatum-DSC was reduced in patients, we controlled for the ROI-based approach with a voxel-wise t-test between patients and healthy controls within the sensorimotor-striatum-ROI, which revealed the dorsal left putamen as peak of reduced DSC (x=-22, y=-4, z=6), although this result did not survive correction for multiple testing. We found no significant associations between sensorimotor-striatum-DSC and CPZ (r=0.09, p=0.70) and FD (r=-0.22, p=0.36), indicating the sensorimotor-striatum-DSC reduction not to be confounded by current medication or head motion.

We found that averaged ASM-thalamic-iFC was increased in patients. This result was supported by voxel-wise two-sample t-test, which revealed hyperconnectivity between ASM and thalamus, peaking in the pulvinar (x=10, y=-24, z=4; cluster-level corrected p_FWE_=0.05 for voxel-wise p=0.001). In patients, there was no significant association between ASM-thalamic-iFC and CPZ (r=0.37, p=0.12) or FD (r=0.12, p=0.61), as shown by Pearson’s correlation analyses, suggesting that ASM-thalamic-iFC results were not confounded by current medication or head motion.

##### *Sensorimotor-striatum-DSC is positively associated with ASM-thalamic-iFC in patients, but decreased sensorimotor-striatum-DSC does not mediate ASM-thalamic-hyperconnectivity*

##### *Associations between sensorimotor-striatum-DSC and ASM-thalamic-iFC using alternative thalamic ROIs*

To control for our choice of thalamic ROI, we tested for associations between sensorimotor-striatum-DSC and ASM-thalamic-iFC in patients by using anatomically defined thalamic nuclei instead of the hyperconnectivity cluster from our previous study (Avram et al., 2018). We found similar results, specifically, sensorimotor-striatum-DSC significantly correlated with ASM-ventrolateral-thalamus-iFC (r=0.66, p=0.004) and with ASM-pulvinar-iFC (r=0.49, p=0.03) (Table S2). We conclude that our finding of associated sensorimotor-striatum-DSC and ASM-thlalamic-iFC was not confounded by our choice of thalamic ROI.

To control for specificity of sensorimotor-striatum-DSC being relevant for ASM-thalamic-iFC, we also tested for associations with additional striatal subdivisions’ DSC and whole striatum DSC. We found associative-striatum-DSC to show a trend association with ASM-thalamic-iFC (r=0.41, p=0.06), but no correlations were found with limbic-striatum-DSC (r=0.28, p=0.15) or whole striatal-DSC (r=0.29, p=0.14). These findings suggest ASM-thalamic-iFC to be specifically associated with the sensorimotor-striatum-DSC.

##### *Mediation analyses for additional striatal subdivisions’ DSC*

Sensorimotor-striatum-DSC did not mediate the effect of schizophrenia on ASM-thalamic-iFC. We further investigated whether other striatal subdivisions might have such an effect. For whole striatum-DSC, mediation analysis revealed that the indirect effect was not significant (95% confidence interval [-0.019 0.01] included zero). Similar, for limbic-striatum-DSC, mediation analysis revealed that the indirect effect was not significant (95% confidence interval [-0.013 0.004] included zero). Finally, for associative-striatum-DSC, mediation analysis revealed that the indirect effect was also not significant (95% confidence interval [-0.02 0.01] included zero). These findings indicate that striatal-DSC does not directly contribute to ASM-thalamic-hyperconnectivity in patients.

##### *Patients’ cognitive difficulties do not link with the ASM-centered-system*

##### *Control for the influence of alternative thalamic ROIs on the association between ASM-thalamic-iFC and BACS scores*

To test whether our choice of thalamic ROI might have an effect on the association between ASM-thalamic-iFC and BACS scores in patients, we repeated the partial correlation analysis using the ventrolateral nucleus and pulvinar, respectively, instead of the hyperconnectivity cluster from our previous study.^10^ No correlations were found for ASM-ventrolateral-thalamus-iFC (r=0.10, p=0.35) and ASM-pulvinar-iFC (r=0.13, p=0.31), suggesting a lack of association between ASM-thalamic-iFC and BACS scores in patients, independent of thalamic ROI. See Table S2 for correlations in healthy controls.

#### Relationships between SAL- and ASM-centered-systems

##### *Sensorimotor-striatum-DSC mediates the group difference in associative-striatum-DSC*

After demonstrating that associative-striatum-DSC mediates the effect of schizophrenia on sensorimotor-striatum-DSC, we also tested whether the opposite holds, namely whether sensorimotor-striatum-DSC mediates the effect of schizophrenia on associative-striatum-DSC. The mediation analysis revealed a significant indirect effect of group on averaged associative-striatum-DSC values via sensorimotor-striatum-DSC (ab=-0.001±0.0007; 95% confidence interval [-0.002 -0.0001] did not include zero). These results indicate that not only dorsomedial striatal-DSC modulated dorsolateral striatal-DSC, but also that dorsolateral striatal-DSC modulated dorsomedial striatal-DSC.

##### *ASM-thalamic-iFC does not mediate the group difference in SAL-thalamic-iFC*

After establishing that SAL-thalamic-iFC mediates the group effect on ASM-thalamic-iFC, we also tested whether the opposite holds, namely whether ASM-thalamic-iFC mediates the effect of schizophrenia on SAL-thalamic-iFC. The mediation was not significant (95% confidence interval [-0.008 0.025] included zero), indicating an anterior-to-posterior effect concerning cortico-thalamic dysconnectivity.

### **Supplementary Discussion**

#### SAL-centered-system changes contribute to ASM-centered-system changes in schizophrenia

##### *Sensorimotor-striatum-DSC mediates the group effect on associative-striatum-DSC*

Our results showed that aberrant sensorimotor-striatum-DSC mediates aberrant associative-striatum-DSC in patients. This result indicated that the dorsolateral striatum is also relevant for the dorsomedial striatum dopamine function. Although this might seem contrary to the ascending spiral concept, recent animal research has demonstrated that associative-striatum and sensorimotor-striatum have parallel, reciprocal connections with the middle and lateral substantia nigra pars compacta, respectively, but that sensorimotor-stratum neurons also strongly project to associative-striatum projecting areas in the substantia nigra pars compacta, which indicates a novel lateral to medial information flow.^13^ Furthermore, beyond the three traditional functional subdivisions of the striatum (limbic, associative, and sensorimotor), a ‘new functional’ region has been recently identified, consisting mainly of the caudate tail, which receives cortical input from several cortical areas, including the auditory cortex.^14^ Studies investigating basal ganglia circuits for reward value-guided behavior have identified parallel circuits passing through the caudate head – voluntary behavior – and caudate tail – automatic behavior - consisting of separate dopaminergic projections, which however, have been hypothesized to interact, so that an automatic process can guide a voluntary action, or a voluntary action can select one of several automatic processes.^15, 16^ Although it is not yet clear where such an interaction could take place, the superior colliculus has been proposed as one possibility, as this region is targeted by both pathways.^16^ To sum up, there is sufficient evidence to suggest that not only the associative-striatum can influence the sensorimotor-striatum but also the other way around.

#### Model: Aberrant associative-striatum-DSC contributes to cortico-thalamic-dysconnectivity

##### *SAL-centered-system and cognitive difficulties*

After investigating the relations between associative-striatum-DSC and SAL-thalamic-iFC, we tested whether associative-striatum-DSC, and SAL-thalamic-iFC, respectively, were linked to cognitive difficulties in patients. We found associative-striatum-DSC to correlate positively with BACS scores, in other words, the lower the associative-striatum-DSC the lower the patients’ performance on BACS (Figure 1, Panel E). Several studies have found a link between associative-striatum-DSC and cognitive functions,^17, 18^ and specifically DSC of the dorsal caudate nucleus (the main part of the associative-striatum) has been implicated in modulating cognitive flexibility in healthy subjects (for a review see^19^). Together, these findings clearly indicate a role of associative-striatum-DSC in cognitive processing, probably based on dopaminergic modulation of ‘cognitive’ networks (see below).

We found SAL-thalamic-iFC to correlate at-trend-to-significance with BACS scores in patients, also suggesting a similar effect: the lower the SAL-thalamic-iFC, the worse the patients’ cognitive performance on BACS (Figure 1, Panel F). This finding is also in line with the literature, as SAL has a central role in the detection of behaviorally relevant stimuli and flexible control of goal-directed behavior,^20^ which make it relevant for cognitive control and working memory.^21^ Regarding patients with schizophrenia, studies have found SAL-thalamic-hypoconnectivity to link with cognitive difficulties.^10, 22^ Furthermore, our results suggest that the mediodorsal thalamus likely drives the association between SAL-thalamic-iFC and BACS scores in patients. This finding is supported by the role the mediodorsal nucleus has in cognition in general, mainly based on dense reciprocal connections with prefrontal areas,^23^ and imaging studies demonstrating decreased mediodorsal activation during tasks evaluating cognitive performance.^24, 25^ Moreover, altered brain activity between the mediodorsal nucleus and the prefrontal cortex has also been reported consistently.^25-27^

Taken together, these results indicate the SAL-centered-system’s involvement in cognitive processing, and demonstrate the relevance of altered dopaminergic signalling and altered iFC within the system for cognitive difficulties in patients with schizophrenia (for a review see Peters and colleagues^28^). Such a role is supported by animal and human imaging studies that suggest striatal dopamine signalling to regulate the activity in cortico-striato-pallido-thalamo-cortical circuits during cognitive processing,^29-31^ possibly by modulating glutamatergic signalling in synapses both at the fronto-striatal and thalamo-striatal levels.^32, 33^

#### Strengths and limitations

##### *Limitations regarding DSC-measure*

Our participants did not take premedication such as entacapone and carbidopa before the PET scan. This pretreatment, which blocks formation of peripheral ^18^F-DOPA-metabolites, increases signal-to-noise-ratio of ^18^F-DOPA-based measures.^34^ Therefore, some effects might not be visible in our study. For further details see Avram and colleagues.^12^

### **Supplementary Tables**

#### Table S1. Antipsychotics and additional psychotropic medication

| **Antipsychotic medication** | **Number of patients (N = 19)** |
| --- | --- |
| Amisulpride | 2 |
| Aripiprazole | 6 |
| Flupentixol | 3 |
| Olanzapine | 7 |
| Paliperidone | 2 |
| Perazine | 1 |
| Perphenazine | 1 |
| Pipamperone | 1 |
| Quetiapine | 4 |
| Risperidone | 1 |
| **Adjunctive psychotropic medication** |  |
| Biperiden | 2 |
| Bupropion | 1 |
| Citalopram | 1 |
| Duloxetine | 1 |
| Escitalopram | 1 |
| Lithium | 1 |
| Methylphenidate | 1 |
| Sertraline | 1 |
| Trimipramine | 1 |
| Valproate | 1 |
| Venlafaxine | 1 |
| Zolpidem | 1 |

#### Table S2. Associations between cortico-thalamic iFC and striatal-DSC: control analyses

| **Partial correlation analyses** | | | | | |
| --- | --- | --- | --- | --- | --- |
|  | **AST-DSC** | **SMST-DSC** | **LST-DSC** | **Striatum DSC** | **BACS** |
| ***Healthy Controls*** |  | | | | |
| **SAL-MD iFC** | r=0.05  p=0.41 | r=0.06  p=0.41 | r=-0.01  p=0.47 | r=-0.04  p=0.44 | r=0.25  p=0.16 |
| **SAL-VA**  **iFC** | r=0.14  p=0.30 | r=0.35  p=0.08 | r=0.32  p=0.11 | r=0.34  p=0.09 | r=0.40  p=0.058 |
| **ASM-VL iFC** | r=-0.002  p=0.49 | r=-0.22  p=0.20 | r=0.31  p=0.12 | r=0.004  p=0.49 | r=-0.10  p=0.34 |
| **ASM-PUL**  **iFC** | r=-0.23  p=0.19 | r=-0.26  p=0.15 | r=-0.05  p=0.41 | r=-0.24  p=0.18 | r=-0.08  p=0.37 |
| ***Patients with SCZ*** |  | | | | |
| **SAL-MD iFC** | r=0.53  p=0.02* | r=0.54  p=0.01* | r=0.26  p=0.17 | r=0.16  p=0.27 | r=0.44  p=0.05* |
| **SAL-VA**  **iFC** | r=0.41  p=0.06 | r=0.40  p=0.06 | r=0.29  p=0.14 | r=0.09  p=0.36 | r=0.27  p=0.16 |
| **ASM-VL iFC** | r=0.50  p=0.02* | r=0.66  p=0.004* | r=0.30  p=0.13 | r=0.40  p=0.06 | r=0.10  p=0.35 |
| **ASM-PUL**  **iFC** | r=0.32  p=0.11 | r=0.49  p=0.03* | r=0.13  p=0.31 | r=0.20  p=0.23 | r=0.13  p=0.31 |

Partial correlation analyses between cortico-thalamic-iFC and DSC (dopamine synthesis capacity) with age, sex, FD (framewise displacement) and for the patient group also CPZ (chlorpromazine equivalents) as covariates of no interest. Abbreviations: SAL – salience network, ASM – auditory sensorimotor network, SCZ – schizophrenia, AST – associative striatum, LST – limbic striatum, SMST – sensorimotor striatum, iFC – intrinsic functional connectivity, MD – mediodorsal nucleus, VA – ventral anterior nucleus, VL – ventrolateral nucleus, PUL - pulvinar. Significant correlations are depicted with *.

### **Supplementary Figures**

#### Figure S1. SPMs, one-sample t-tests, for striatal-DSC and SAL-thalamic/ASM-thalamic-iFC

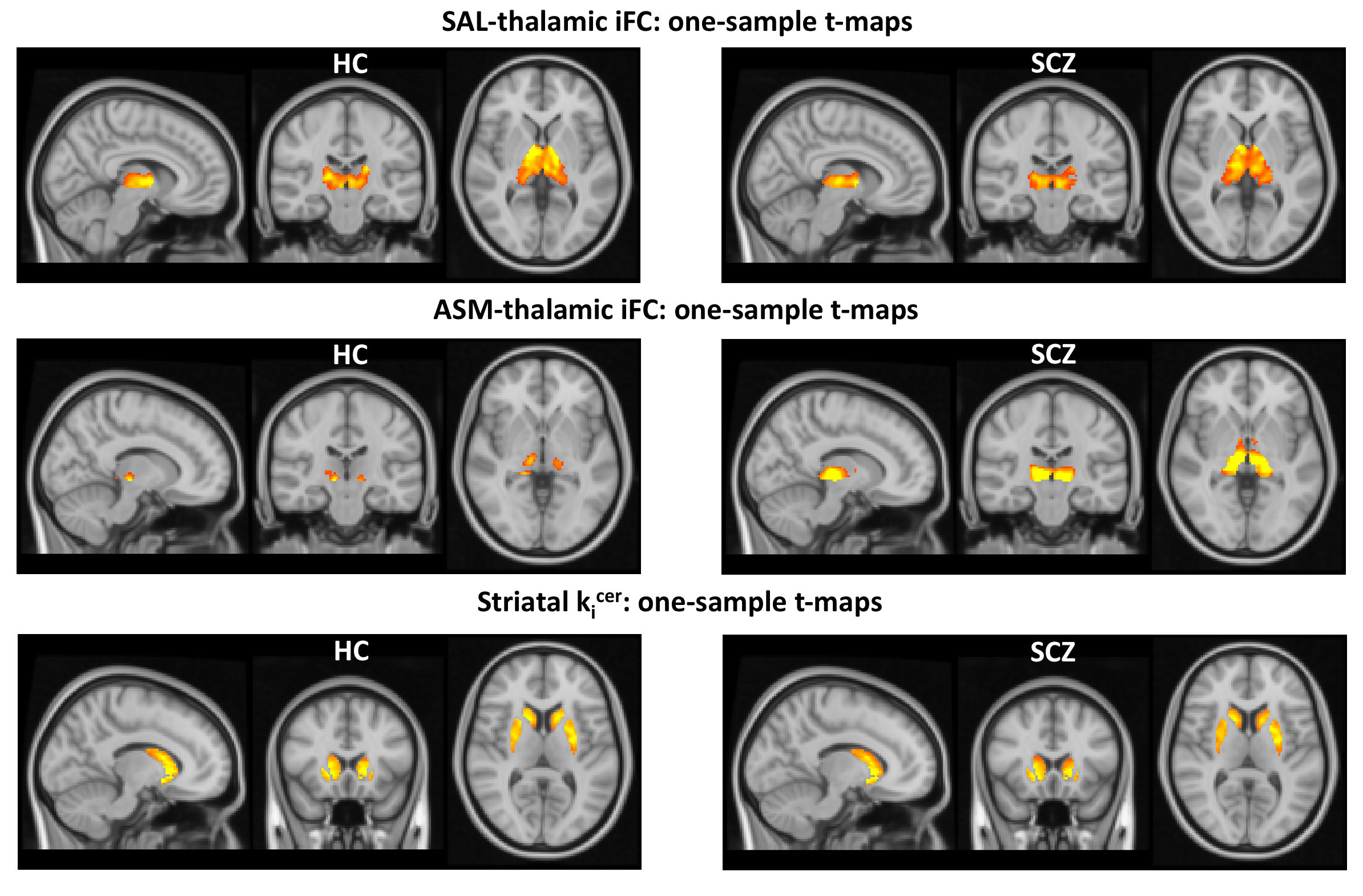

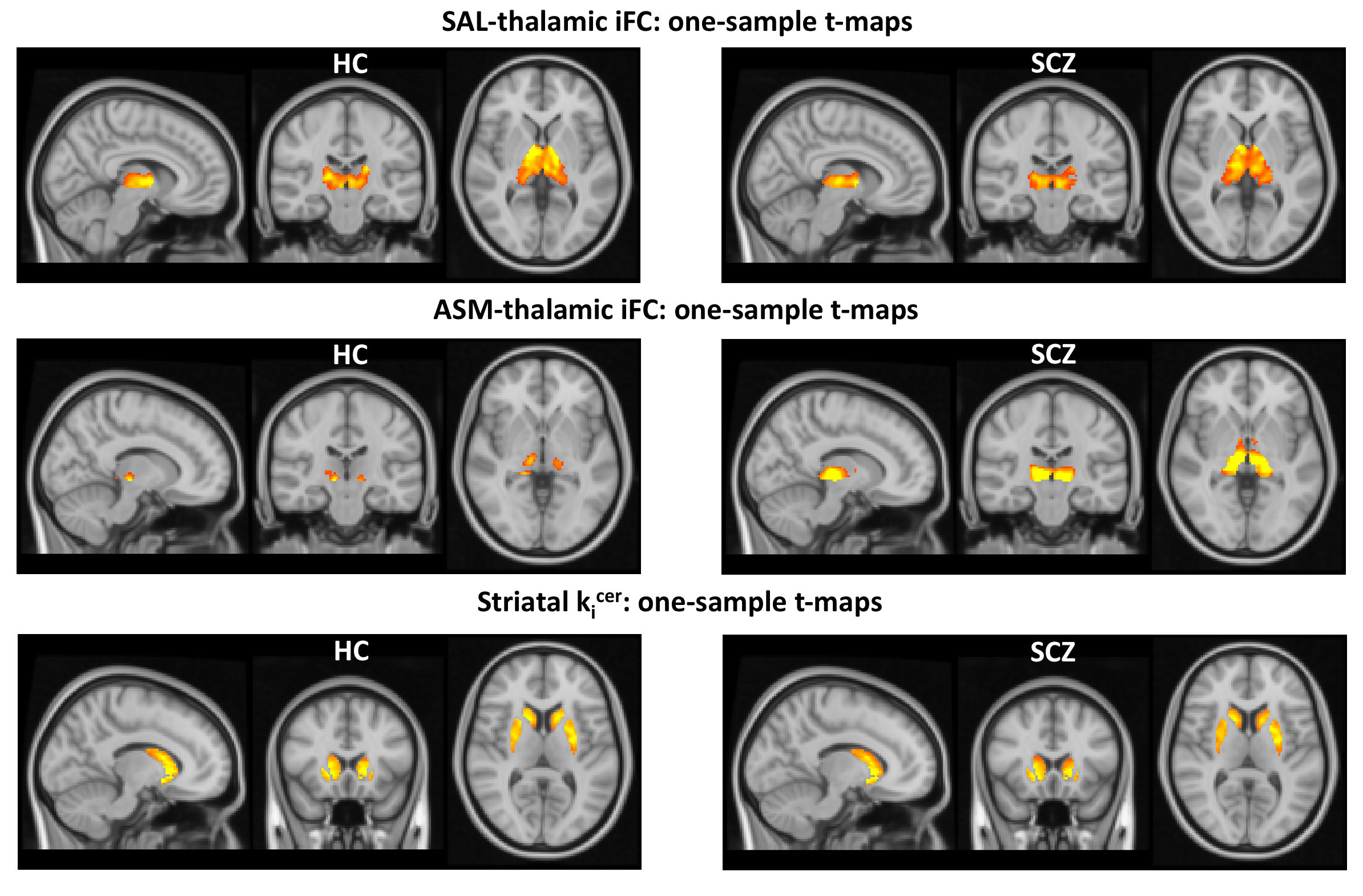

One-sample t-maps for healthy controls (HC) and patients with schizophrenia (SCZ), cluster level corrected p_FWE_<0.05, for striatal dopamine synthesis capacity k_i_^cer^, SAL-thalamic-iFC, and ASM-thalamic-iFC. Abbreviations: SAL – salience network, ASM – auditory-sensorimotor network, iFC – intrinsic functional connectivity, k_i_^cer^ – reflects dopamine synthesis capacity.

#### Figure S2. Overlap of anatomically defined thalamic nuclei with hypo- and hyperconnectivity clusters from our previous study (Avram et al., 2018)

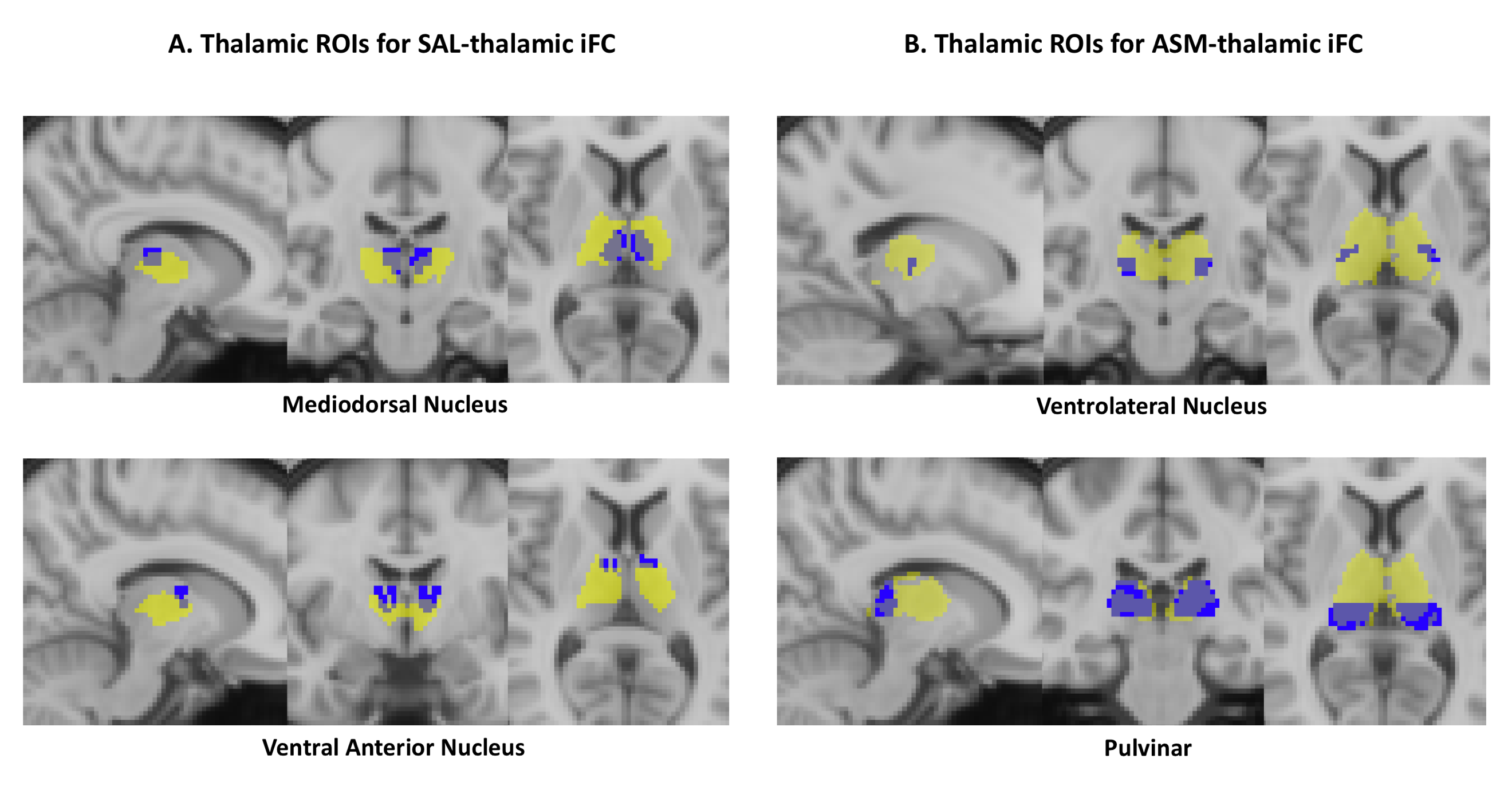

Overlap of hypo-/hyperconnectivity clusters (in transparent yellow) on anatomically defined thalamic nuclei (in blue). Abbreviations: ROI – region of interest, SAL – salience network, ASM – auditory-sensorimotor network, iFC – intrinsic functional connectivity.

#### Figure S3. Relationships between SAL-centered and ASM-centered-systems

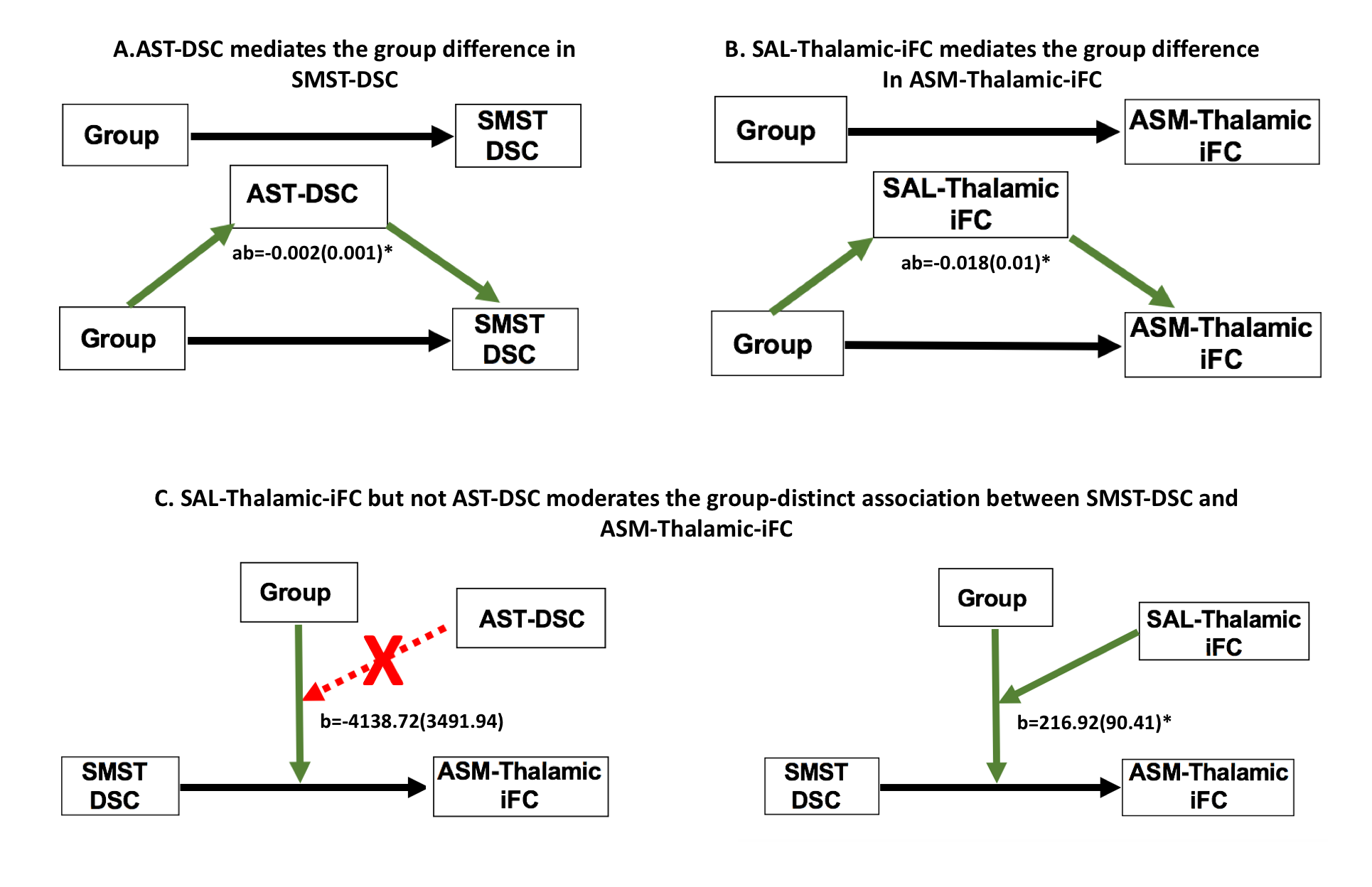

**A.** Associative-striatum-DSC mediates the group difference in sensorimotor-striatum-DSC. Top: total effect – ‘path c’, reflecting the ‘simple’ association between the factor ‘group’ and sensorimotor-striatum-DSC. Bottom: ‘path a’, reflecting effect of causal variable (group) on mediator (associative-striatum-DSC); ‘path b’, reflecting the effect of the mediator (associative-striatum-DSC) on the outcome variable (sensorimotor-striatum-DSC); ‘path ab’ – reflecting the indirect effect (mediation was significant), regression coefficient and standard error are shown; ‘path c’’, reflecting the direct effect of the causal variable (group) on the outcome variable (sensorimotor-striatum-DSC). **B.** SAL-thalamic-iFC mediates the group difference in ASM-thalamic-iFC. Top: total effect – ‘path c’, reflecting the ‘simple’ association between the factor ‘group’ and ASM-thalamic iFC. Bottom: ‘path a’, reflecting effect of causal variable (group) on mediator (SAL-thalamic-iFC); ‘path b’, reflecting the effect of the mediator (SAL-thalamic-iFC) on the outcome variable (ASM-thalamic-iFC); ‘path ab’ – reflecting the indirect effect (mediation was significant), regression coefficient and standard error are shown; ‘path c’’, reflecting the direct effect of the causal variable (group) on the outcome variable (ASM-thalamic-iFC). **C.** SAL-thalamic-iFC but not associative-striatum-DSC moderates the group-distinct association between sensorimotor-striatum-DSC and ASM-thalamic-iFC. *Left*: three-way-interaction revealed that associative-striatum-DSC does not moderate the group-distinct association between sensorimotor-striatum-DSC and ASM-thalamic-iFC. Beta coefficient and standard error are shown for the interaction between associative-striatum-DSC*group*sensorimotor-striatum-DSC (p=0.15). *Right*: three-way-interaction revealed that SAL-thalamic-iFC moderates the group-distinct association between sensorimotor-striatum-DSC and ASM-thalamic-iFC. Beta coefficient and standard error are shown for the interaction between SAL-thalamic-iFC*group*sensorimotor-striatum-DSC (p=0.02). *Abbreviations*: SAL – salience network, ASM – auditory-sensorimotor network, AST- associative-striatum, SMST – sensorimotor-striatum, DSC – dopamine synthesis capacity, iFC – intrinsic functional connectivity.
